# A trimeric hydrophobic zipper mediates the intramembrane assembly of SARS-CoV-2 spike

**DOI:** 10.1101/2021.04.09.439203

**Authors:** Qingshan Fu, James J. Chou

## Abstract

The S protein of the SARS-CoV-2 is a Type I membrane protein that mediates membrane fusion and viral entry. A vast amount of structural information is available for the ectodomain of S, a primary target by the host immune system, but much less is known regarding its transmembrane domain (TMD) and its membrane-proximal regions. Here, we determined the nuclear magnetic resonance (NMR) structure of the S protein TMD in bicelles that closely mimic a lipid bilayer. The TMD structure is a transmembrane α-helix (TMH) trimer that assembles spontaneously in membrane. The trimer structure shows an extensive hydrophobic core along the 3-fold axis that resembles that of a trimeric leucine/isoleucine zipper, but with tetrad, not heptad, repeat. The trimeric core is strong in bicelles, resisting hydrogen-deuterium exchange for weeks. Although highly stable, structural guided mutagenesis identified single mutations that can completely dissociate the TMD trimer. Multiple studies have shown that the membrane anchor of viral fusion protein can form highly specific oligomers, but the exact function of these oligomers remain unclear. Our findings should guide future experiments to address the above question for SARS coronaviruses.

---

The SARS-CoV-2 virion is decorated with a large number of membrane-anchored spike proteins (S) responsible for target recognition, membrane fusion, and virus entry^1-2^; it is also the dominant antigen on the virion surface used for vaccine development^3^. The full-length S is a Type I membrane protein that is first expressed as a precursor that trimerizes (S_3_) and then cleaved into two fragments ((S1/S2)_3_): the receptor-binding fragment S1 and the fusion fragment S2^4^.

The processed (S1/S2)_3_ comprises the crown-shaped ectodomain that contains the receptor binding domain (RBD), a transmembrane domain (TMD), and a cytoplasmic tail (CT) (Fig. 1a). Since the availability of the SARS-CoV-2 genetic code in January of 2020, structural biology of the SARS-CoV-2 spike has progressed at a lightning speed owing to cryo-electron microscopy (cryo-EM), i.e., over 26 structures of the S1/S2 ectodomain have been published, most of them covering residues 14–1162 (Table S1). But as has been the case for the spike proteins of many enveloped viruses, the membrane region of the coronavirus spike remains unknown. In a cryo-EM study that thus far provided the most complete view of the S protein, structural details could be seen up to the HR2 region of the S2 fragment (Fig. 1a) but the membrane-proximal and transmembrane regions were not resolved^5^.

**Figure 1.**
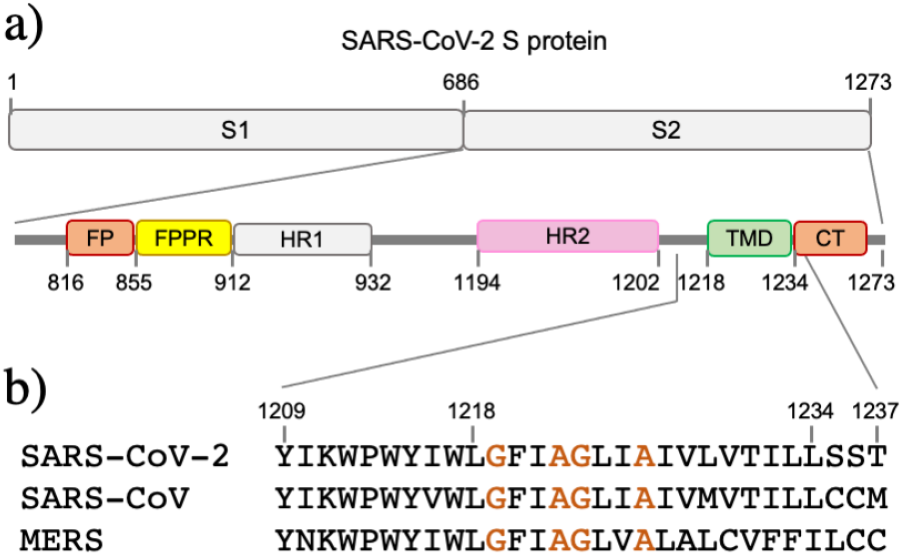
Sequence arrangement of the membrane-interacting regions of SARS-CoV-2 S. (**a**) Overall domain organization of S2. (**b**) Sequence alignment of the TMDs of S2 of SARS-CoV-2 (QII57161.1), SARS-CoV (AAS75868.1), and MERS (QDI73610.1).

Previous studies on SARS-CoV, however, suggest that the S TMD has important functional roles other than membrane anchoring. One study showed that swapping the TMD of SARS-CoV S with that of vesicular stomatitis virus (VSV) G protein resulted in 3-25% activity compared to the wildtype^6^. Another study reported that insertion of a residue in the TMD resulted in a complete block of viral entry^7^. Further, a recent study on SARS-CoV-2 showed that directly fusing the RBD to the TMD could induce trimerization, suggesting the ability of the TMD to trimerize^8^.

In this study, we used NMR to investigate the structural properties of the TMD of SARS-CoV-2 S in bicelles to fill the knowledge gap. We find that the TMD of the S protein forms a strong trimer in bicelles by a previously unknown mode of transmembrane helix (TMH) assembly.

To determine the TMD structure using NMR, we used a S2 fragment (residues 1209-1237; Fig. 1b), derived from a SARS-CoV-2 isolate QII57161.1. This construct, designated S2^1209-1237^, contains a short stretch of the membrane-proximal region (residues 1209-1217) and the TM segment (residues 1218-1234). S2^1209-1237^ was reconstituted in DMPC-DH_6_PC bicelles with *q* = 0.55 (Fig. S1a,b), where *q* = [DMPC]/[DH_6_PC]. At *q* = 0.55, the diameter of the planar bilayer region of the bicelles is ∼46 Å^9^. The bicelle-reconstituted S2^1209-1237^ ran on SDS-PAGE as trimers, whereas unreconstituted peptide migrated as monomers (Fig. S1c). Further, OG-label analysis independently showed that S2^1209-1237^ forms trimers in bicelles (Fig. S1d).

The trimeric S2^1209-1237^ in bicelles generated good NMR spectra (Fig. S2), and its NMR structure was determined using a published protocol^10^, involving 1) construction of a preliminary monomer structure with local nuclear Overhauser effect (NOE) restraints and backbone dihedral angles derived from chemical shift values (using TALOS+^11^), 2) obtaining a unique structural solution (using ExSSO^12^) that satisfies inter-chain NOE restraints derived from mixed isotopically labeled sample (Fig. S3), and 3) refinement of the trimer structure by further assignment of self-consistent NOE restraints (overall procedure in Fig. S4; refinement summary in Table S2).

In bicelles, the TMD of SARS-CoV-2 S protein folds into a regular α-helix (residues 1218-1234) that assembles into a parallel homotrimer (Fig. 2a). Residues 1209-1217 are unstructured in our sample, likely due to N-terminal truncation. The trimeric complex is held together by an extensive hydrophobic core along the 3-fold axis, and the core comprises four layers of hydrophobic interaction involving I1221, I1225, L1229, and L1233, respectively (Fig. 2a). Despite the presence of signature sequences for driving TMH oligomerization such as Gly-xxx-Gly and Ala-xxx-Ala13-14, our TMD structure does not show direct involvement of the glycine or alanine in forming close van der Waals (VDW) contacts. In this regard, the new TMH trimerization mode is different from the known structures that require one or two small amino acids in establishing intimate helix-helix contact, e.g., the G690 for HIV-1 gp41^15^, the G221 for TNFR1^16^, a central proline for Fas^17^, a central alanine for DR5^18^, and the A794 for HSV gB^19^ TMD structures.

**Figure 2.**
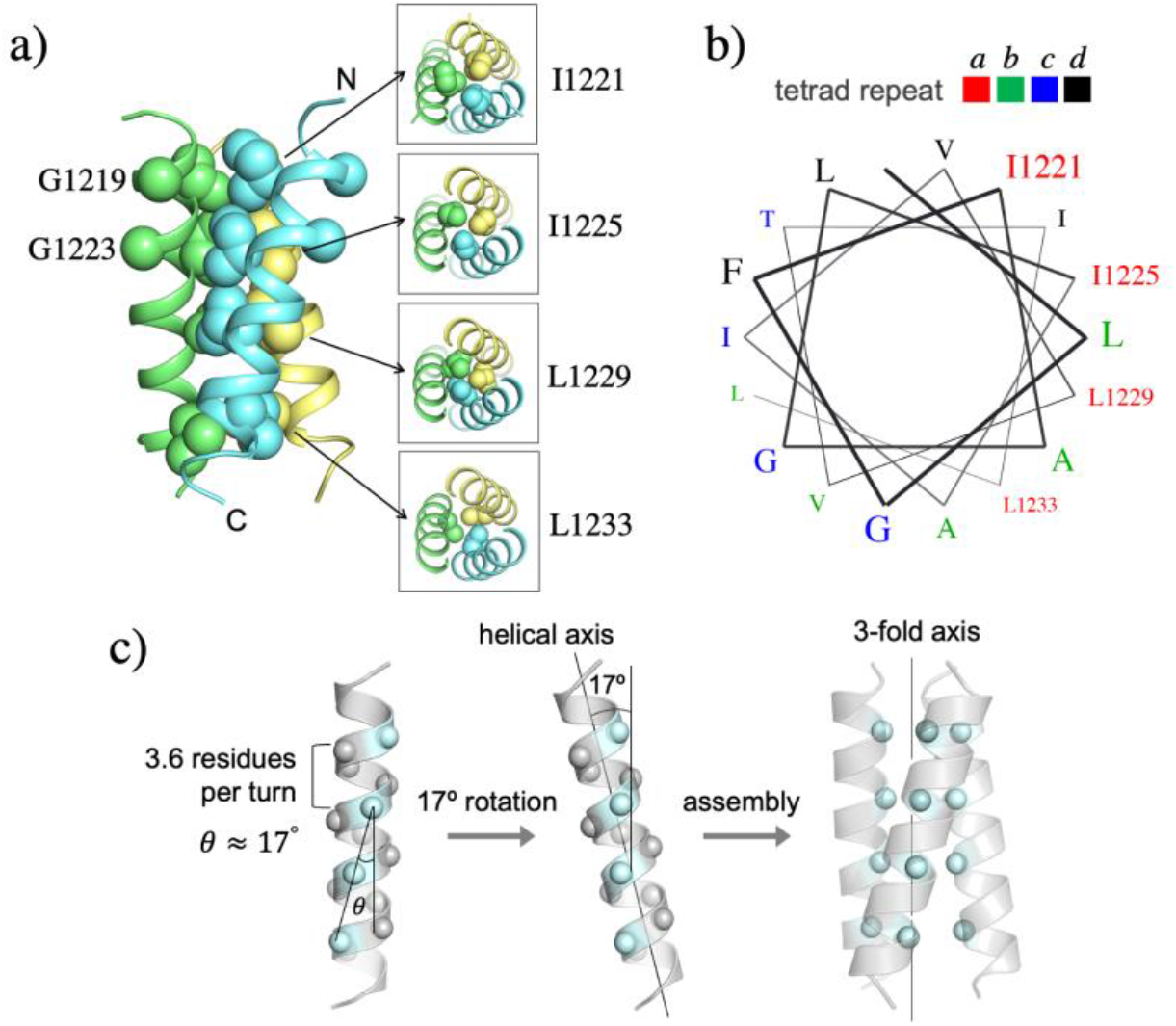
NMR structure of the TMH trimer of SARS-CoV-2 S in DMPC-DH_6_PC bicelle with *q* = 0.55. (**a**) Ribbon representation (left) of the TMH trimer structure with the sidechain heavy atoms of the core residues shown as spheres; the Cα atoms of G1219 and G1223 are also shown as spheres. The sidechain packing at four different levels along the threefold axis is illustrated with sectional top views of the trimer (right). (**b**) Helical wheel representation of an α-helix (3.6 residues per turn) showing that the core hydrophobic residues occupy the position ‘*a*’ of the ‘*abcd*’ tetrad repeat. (**c**) Theoretical analysis of the trimeric hydrophobic zipper with tetrad repeat. The line formed by the Cα atoms of the residues at position ‘*a*’ is tilted by ∼17º relative to the helical axis. Rotating the helix by 17º places the position-*a* residues in line with the 3-fold axis for optimal hydrophobic core formation.

The hydrophobic core of the TMD trimer shows an unusual pattern of tetrad repeat, i.e., I1221, I1225, L1229, and L1233, each occupying position ‘*a*’ of the *abcd* repeat (Fig. 2b), and this is very different from the coiled coil mode of assembly of TMH with heptad repeat^20^. Since each turn of an α-helix consists of 3.6 residues, a 4-residue repeat overshoots the *i* + 4 hydrophobic residues past a helical turn by 40°, diverting the hydrophobic ridge from the 3-fold axis by ∼17° (Fig. 2c). Thus, tilting the TMH by 17° would align the hydrophobic ridges of the three TMHs with the 3-fold axis to allow intimate hydrophobic packing (Fig. 2c). Indeed, the tilt angle in our experimentally determined structure is ∼19°, in close agreement with the theoretical analysis.

To examine the S TMD independently by mutagenesis, we generated seven single mutations – G1219Y, G1223Y, I1221Y, I1225Y, A1226Y, L1229Y, and L1233Y – and evaluated their effect on TMH trimerization (Fig. 3a). Mutating the characteristic glycine/alanine in the Gly^1219^-xxx-Gly^1223^ or Ala^1222^-xxx-Ala^1226^ signature sequence to tyrosine has no effect on TMH trimerization, further supporting the structural conclusion that the relatively conserved glycine and alanine are not directly involved in helix-helix packing. As shown in Fig. 2a, G1219 and G1223 are entirely lipid facing, not expected to participate in interhelical VDW contacts. A1226 is closer to the packing interface but is still not interior enough to be in VDW contact with I1225 or L1229 of the neighboring chain. In contrast, mutating each of the four hydrophobic residues (I1221, I1225, L1229, L1233) that constitute the hydrophobic core to tyrosine all led to severe disruption of the trimer, further consolidating the conclusion that the tetrad repeat of bulky hydrophobic residues is important for the TMH trimerization. Further, the I1225Y or L1229Y mutation almost completely abolished trimerization while the I1221Y and L1233Y near the N- and C-terminal ends of the TMH, respectively, are less disruptive, probably due to increased dynamics of the helix-ends (Fig. S5). This is also consistent with the hydrogen-deuterium (H-D) exchange that the core residues 1225 – 1229 exhibited the lowest *k*_ex_ of all residues (Fig. 3b; Fig. S6). Overall, the oligomeric properties of the seven mutants agree well with the NMR structure.

**Figure 3.**
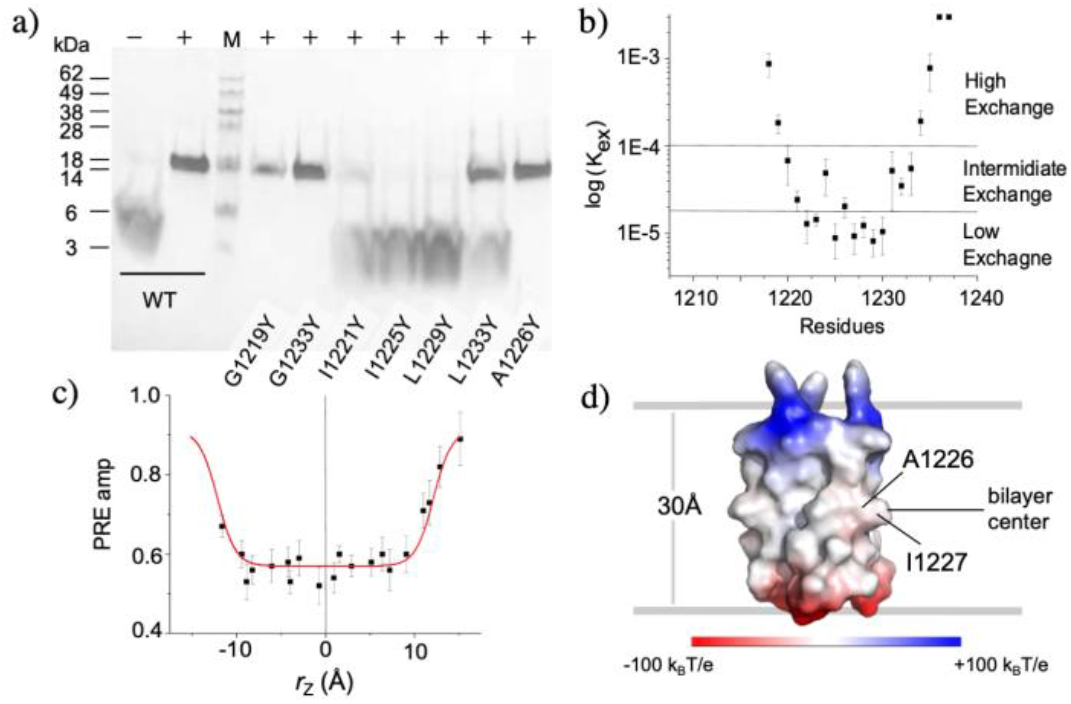
Stability and localization of the TMH trimer of SARS-CoV-2 S in bicelles. (**a**) SDS-PAGE of bicelle-reconstituted S2^1209-1237^ and its mutants for showing the effect of single mutations on trimerization. Samples were run under non-denaturing conditions. ‘−’ and ‘+’ indicate unreconstituted and bicelle reconstituted, respectively. (**b**) Residue-specific amide *k*_ex_ at pH 6.8 determined by H-D exchange measurements. (**c**) *PRE*_amp_ vs *r*_Z_ best fitted to symmetric sigmoidal equation (Eq. S2), where *r*_Z_ = 0 corresponds to the bilayer center. (**d**) Position of the TMH trimer (electrostatic surface representation) relative to the center and boundaries of the planar region of the bicelle.

To determine the membrane partition of the TMD structure, we performed the paramagnetic probe titration (PPT) analysis^10, 21^ using S2^1209-1237^ reconstituted in bicelles with *q* = 0.6. Soluble (Gd-DOTA) and lipophilic (16-DSA) probes were used to provide reciprocal paramagnetic relaxation enhancement (PRE) information (Fig. S7). The analysis of residue-specific PRE amplitude (*PRE*_amp_) in the context of the TMD structure shows that the lipid bilayer center is between A1226 and I1227 (Fig. 3c, d; Fig. S8; Table S3). Further, *PRE*_amp_ reaches the maximal values at about 15Å away from the center on either side, indicating that the bilayer thickness around the TMD trimer is ∼30Å. Thus, the S protein TMD caused substantial thinning of the membrane around it.

We have shown that the TMD of the SARS-CoV-2 fusion protein spontaneously trimerizes in lipid bilayer, and the trimeric assembly is achieved with a previously unknown hydrophobic zipper motif with tetrad repeat, not with the usual suspects of TMH oligomerization motifs containing glycine or alanine. The role of small amino acids in mediating TMH oligomerization has been observed in several Type I/II membrane proteins including glycophorin A^13^, growth factor receptors^22-23^, and receptors in the tumor necrosis factor receptor (TNFR) superfamily^16-18^. The Gly-xxx-Gly is a well-known motif that drives TMH dimerization^13-14, 22, 24^. There have been no reports, however, of the Gly-xxx-Gly involvement in TMH trimerization. In the trimer structure of the HIV-1 Env TMD, which contains a highly conserved Gly-xxx-Gly, only the first glycine is involved in helix-helix packing; the second glycine is lipid facing^15, 25^. SARS-CoV-2 S TMD also contains highly conserved small amino acids (G1219, A1222, G1223, A1226), which we thought initially to be important for TMD oligomerization. But, neither of the glycines and alanines in the trimer structure appears to be important for the hydrophobic core formation. The purpose of the Gly-xxx-Gly motif remains unknown. If the trimer structure presented here represents the prefusion state, a possible role of the glycine motif is in later steps of the fusion mechanism.

Although the TMH is relatively short (∼16 residues), it can have an extensive hydrophobic core with four layers of hydrophobic interaction. This is attributed to the tetrad repeat of hydrophobic residues, as opposed to the heptad repeat in a classic coiled coil structure. Based on the extensive hydrophobic packing, we believe the TMD trimer is stable in membrane. A potential implication is that the TMD trimer is unlikely to dissociate in the membrane unless significant force is applied during the unfolding and refolding steps of the fusion component.

Although functional mutagenesis of the SARS-CoV-2 S TMD has not been reported, a previous study on the SARS-CoV reported that inserting an amino acid between G1201 and F1202 of the S TMD completely blocked viral entry^7^. G1201 and F1202 in SARS-CoV correspond to G1219 and F1220 in SARS-CoV-2, respectively (Fig. 1b). In the context of our TMH trimer structure, such insertion might not disrupt trimerization but could place the tetrad repeat out of register relative to the still unknown membrane-proximal structure and thus prevent proper TMH trimerization.

In conclusion, the TM anchor of the SARS-CoV-2 fusion protein spontaneously trimerizes in the membrane. The trimeric complex is stabilized by an extensive hydrophobic core along the 3-fold axis, formed by the bulky hydrophobic amino acids repeated every four residues. This mode of TMH trimerization is significantly different from the known TMH trimer structures of fusion proteins from other viruses. Strong intramembrane oligomerization appears to be a recurring theme for viral fusion proteins, but its functional roles remain unclear. The reported structure of the TMD of SARS-CoV-2 fusion protein allowed us to identify single mutations that can completely dissociate the trimeric assembly. We believe these mutations are valuable information for guiding future functional experiments for addressing the above question.

## ASSOCIATED CONTENT

### Supporting Information

The Supporting Information is available free of charge on the ACS Publications website at DOI: xxxx. The SI includes Tables S1-S4 and Fig. S1-S9, as well as the description of sample preparation, NMR and biochemical analyses.

## AUTHOR CONTRIBUTION

Q.F., J.J.C. conceived the study; Q.F. designed protein constructs and prepared samples; Q.F., J.J.C. determined the NMR structure and designed mutants for functional study; J.J.C. wrote the paper and Q.F. contributed to editing of the manuscript.

## ACKNOWLEDGEMENT

We thank Bo OuYang and Bing Chen for insightful discussion. This study was supported by NIH grant AI127193 to J.J.C. The NMR data were collected at the MIT-Harvard CMR (supported by NIH grant P41 EB-002026).

---

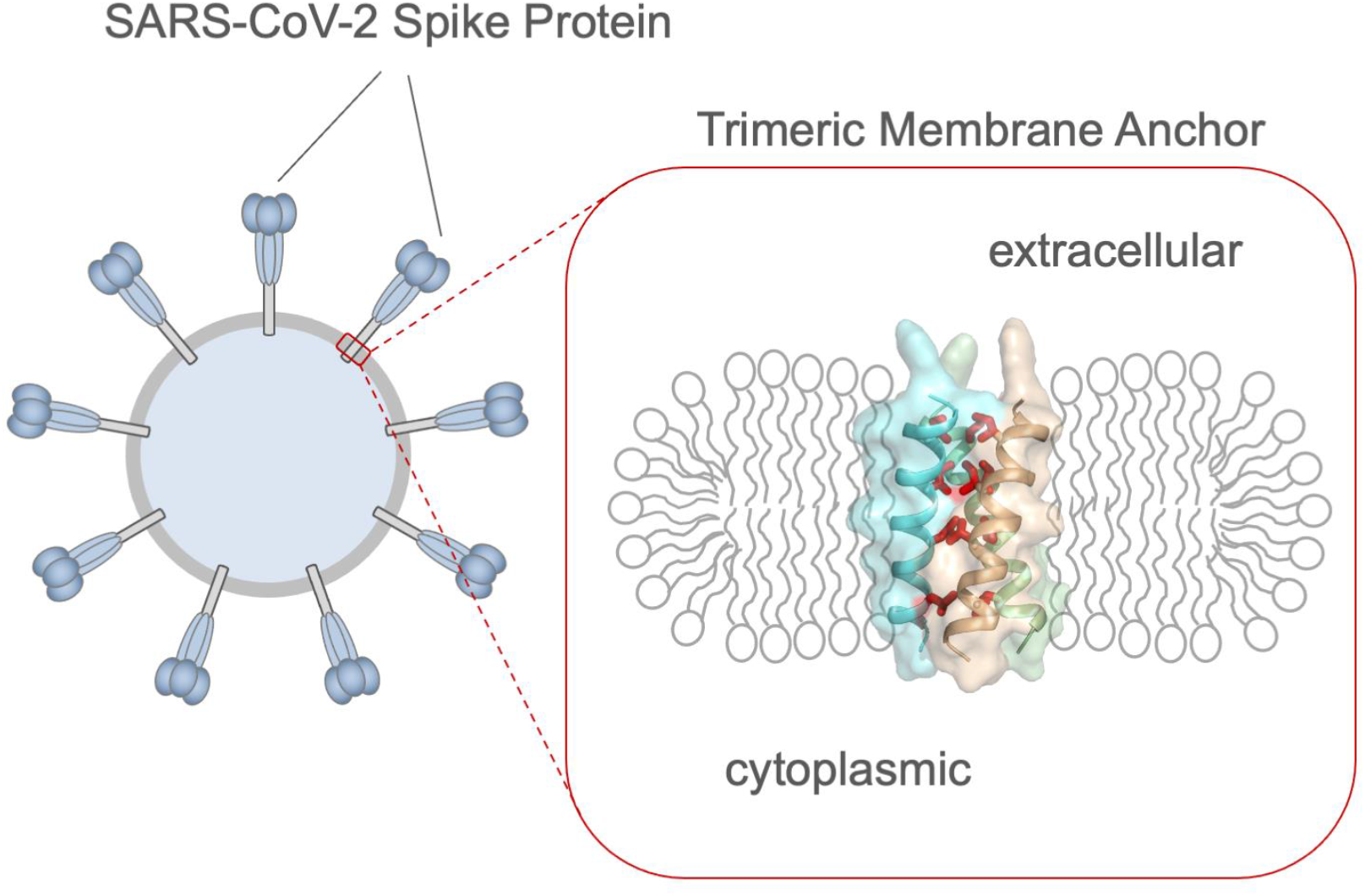

## Notes

### Competing Interest Statement

The authors have declared no competing interest.

## REFERENCES

1. Klein, S., Cortese, M., Winter, S. L., Wachsmuth-Melm, M., Neufeldt, C. J., Cerikan, B., Stanifer, M. L., Boulant, S., Bartenschlager, R., Chlanda, P., SARS-CoV-2 structure and replication characterized by in situ cryo-electron tomography. Nat Commun 2020, 11 (1), 5885.

2. Benton, D. J., Wrobel, A. G., Xu, P., Roustan, C., Martin, S. R., Rosenthal, P. B., Skehel, J. J., Gamblin, S. J., Receptor binding and priming of the spike protein of SARS-CoV-2 for membrane fusion. Nature 2020, 588 (7837), 327–330.

3. Lv, Z., Deng, Y. Q., Ye, Q., Cao, L., Sun, C. Y., Fan, C., Huang, W., Sun, S., Sun, Y., Zhu, L., Chen, Q., Wang, N., Nie, J., Cui, Z., Zhu, D., Shaw, N., Li, X. F., Li, Q., Xie, L., Wang, Y., Rao, Z., Qin, C. F., Wang, X., Structural basis for neutralization of SARS-CoV-2 and SARS-CoV by a potent therapeutic antibody. Science 2020, 369 (6510), 1505–1509.

4. Bosch, B. J., van der Zee, R., de Haan, C. A., Rottier, P. J., The coronavirus spike protein is a class I virus fusion protein: structural and functional characterization of the fusion core complex. J Virol 2003, 77 (16), 8801–11.

5. Cai, Y., Zhang, J., Xiao, T., Peng, H., Sterling, S. M., Walsh, R. M., Rawson, S., Rits-Volloch, S., Chen, B., Distinct conformational states of SARS-CoV-2 spike protein. bioRxiv 2020.

6. Broer, R., Boson, B., Spaan, W., Cosset, F. L., Corver, J., Important role for the transmembrane domain of severe acute respiratory syndrome coronavirus spike protein during entry. J Virol 2006, 80 (3), 1302–10.

7. Corver, J., Broer, R., van Kasteren, P., Spaan, W., Mutagenesis of the transmembrane domain of the SARS coronavirus spike glycoprotein: refinement of the requirements for SARS coronavirus cell entry. Virol J 2009, 6, 230.

8. Azad, T., Singaravelu, R., Crupi, M. J. F., Jamieson, T., Dave, J., Brown, E. E. F., Rezaei, R., Taha, Z., Boulton, S., Martin, N. T., Surendran, A., Poutou, J., Ghahremani, M., Nouri, K., Whelan, J. T., Duong, J., Tucker, S., Diallo, J. S., Bell, J. C., Ilkow, C. S., Implications for SARS-CoV-2 Vaccine Design: Fusion of Spike Glycoprotein Transmembrane Domain to Receptor-Binding Domain Induces Trimerization. Membranes (Basel) 2020, 10 (9).

9. Sanders, C. R., 2nd; Schwonek, J. P., Characterization of magnetically orientable bilayers in mixtures of dihexanoylphosphatidylcholine and dimyristoylphosphatidylcholine by solid-state NMR. Biochemistry 1992, 31 (37), 8898–905.

10. Fu, Q., Piai, A., Chen, W., Xia, K., Chou, J. J., Structure determination protocol for transmembrane domain oligomers. Nat Protoc 2019, 14 (8), 2483–2520.

11. Shen, Y., Delaglio, F., Cornilescu, G., Bax, A., TALOS+: a hybrid method for predicting protein backbone torsion angles from NMR chemical shifts. Journal of Biomolecular NMR 2009, 44 (4), 213–23.

12. Yang, J., Piai, A., Shen, H. B., Chou, J. J., An Exhaustive Search Algorithm to Aid NMR-Based Structure Determination of Rotationally Symmetric Transmembrane Oligomers. Sci Rep 2017, 7 (1), 17373.

13. MacKenzie, K. R., Prestegard, J. H., Engelman, D. M., A transmembrane helix dimer: structure and implications. Science 1997, 276 (5309), 131–3.

14. Trenker, R., Call, M. E., Call, M. J., Crystal Structure of the Glycophorin A Transmembrane Dimer in Lipidic Cubic Phase. Journal of the American Chemical Society 2015, 137 (50), 15676–9.

15. Dev, J., Park, D., Fu, Q., Chen, J., Ha, H. J., Ghantous, F., Herrmann, T., Chang, W., Liu, Z., Frey, G., Seaman, M. S., Chen, B., Chou, J. J., Structural basis for membrane anchoring of HIV-1 envelope spike. Science 2016, 353 (6295), 172–175.

16. Zhao, L., Fu, Q., Pan, L., Piai, A., Chou, J. J., The Diversity and Similarity of Transmembrane Trimerization of TNF Receptors. Front Cell Dev Biol 2020, 8, 569684.

17. Fu, Q., Fu, T. M., Cruz, A. C., Sengupta, P., Thomas, S. K., Wang, S., Siegel, R. M., Wu, H., Chou, J. J., Structural Basis and Functional Role of Intramembrane Trimerization of the Fas/CD95 Death Receptor. Mol Cell 2016, 61 (4), 602–613.

18. Pan, L., Fu, T. M., Zhao, W., Zhao, L., Chen, W., Qiu, C., Liu, W., Liu, Z., Piai, A., Fu, Q., Chen, S., Wu, H., Chou, J. J., Higher-Order Clustering of the Transmembrane Anchor of DR5 Drives Signaling. Cell 2019, 176 (6), 1477–1489 e14.

19. Cooper, R. S., Georgieva, E. R., Borbat, P. P., Freed, J. H., Heldwein, E. E., Structural basis for membrane anchoring and fusion regulation of the herpes simplex virus fusogen gB. Nat Struct Mol Biol 2018, 25 (5), 416–424.

20. Rout, A. K., Strub, M. P., Piszczek, G., Tjandra, N., Structure of transmembrane domain of lysosome-associated membrane protein type 2a (LAMP-2A) reveals key features for substrate specificity in chaperone-mediated autophagy. The Journal of biological chemistry 2014, 289 (51), 35111–23.

21. Piai, A., Fu, Q., Dev, J., Chou, J. J., Optimal Bicelle Size q for Solution NMR Studies of the Protein Transmembrane Partition. Chemistry 2017, 23 (6), 1361–1367.

22. Bocharov, E. V., Mineev, K. S., Volynsky, P. E., Ermolyuk, Y. S., Tkach, E. N., Sobol, A. G., Chupin, V. V., Kirpichnikov, M. P., Efremov, R. G., Arseniev, A. S., Spatial structure of the dimeric transmembrane domain of the growth factor receptor ErbB2 presumably corresponding to the receptor active state. The Journal of biological chemistry 2008, 283 (11), 6950–6.

23. Endres, N. F., Das, R., Smith, A. W., Arkhipov, A., Kovacs, E., Huang, Y., Pelton, J. G., Shan, Y., Shaw, D. E., Wemmer, D. E., Groves, J. T., Kuriyan, J., Conformational coupling across the plasma membrane in activation of the EGF receptor. Cell 2013, 152 (3), 543–56.

24. Chen, W., Gamache, E., Rosenman, D. J., Xie, J., Lopez, M. M., Li, Y. M., Wang, C., Familial Alzheimer’s mutations within APPTM increase Abeta42 production by enhancing accessibility of epsilon-cleavage site. Nat Commun 2014, 5, 3037.

25. Chen, B., Chou, J. J., Structure of the transmembrane domain of HIV-1 envelope glycoprotein. FEBS J 2017, 284 (8), 1171–1177.

