## Supplementary material for "A trimeric hydrophobic zipper mediates the intramembrane assembly of SARS-CoV-2 spike": SI: SI-for-pub.pdf

Qingshan Fu and James J. Chou

Department of Biological Chemistry and Molecular Pharmacology, Harvard Medical School, Boston, Massachusetts 02115, USA

### Materials and Methods

Protein expression and purification

DNA of a fragment of SARS-CoV-2 S2 (isolate QII57161.1) containing residues 1209-1236 was synthesized by GenScript (Piscataway, NJ). The expression construct, designated $S2^{1209\text{-}1237}$, was created as a N terminal trpLE fusion in the plasmid pMM-LR6 following a procedure described previously[1]. Mutant constructs were generated by standard PCR mutation protocols and confirmed by DNA sequencing. Transformed *E. coli* strain BL21 (DE3) cells were grown in M9 minimal media supplemented with stable isotopes. Cultures were grown at 37°C to reach an absorbance of 0.6~0.8 at 600 nm and cooled to 22°C before induction with 150 μM isopropyl β-D-thiogalatopyranoside at 22° C for overnight (12 hours). For completely deuterated proteins, bacterial culture in 1 ml LB medium was spun down and the cells were adapted in 99.8% $D_2O$ medium (100 ml, with deuterated glucose) over night. Then collected cells were grown in 99.8% $D_2O$ (4 L) (Sigma Aldrich, St. Louis, MO) with deuterated glucose (99.8% deuterium, Cambridge Isotope Laboratories, Tewksbury, MA). The expressed fusion protein was extracted in a denaturing buffer (1% Triton X-100, 6 M guanidine hydrochloride, 50 mM Tris, pH 8.0, and 200 mM NaCl), purified with a Ni-NTA affinity column, cleaved by cyanogen bromide (CNBr) to separate trpLE and $S2^{1209\text{-}1237}$. The final product was purified RP-HPLC in a Zorbax SB-C3 column (Agilent Technologies, Santa Clara, CA) with a gradient from 95% $dH_2O$, 5% isopropanol (IPA), 0.1% trifluoroacetic acid (TFA) (buffer A) to 75% IPA, 25% acetonitrile, 0.1% TFA (buffer B). Fractions containing pure $S2^{1209\text{-}1237}$ was confirmed by MALDI-TOF mass spectrometry and SDS-PAGE.

Reconstitution

Approximately 2 mg of lyophilized $S2^{1209\text{-}1237}$ was mixed with 10 mg 1,2-dimyristoyl-sn-Glycero-3-Phosphocholine (DMPC; protonated or deuterated from Avanti Ploar Lipids, Alabaster, AL) and dissolved in 1,1,1,3,3,3-hexafluoro-2-propanol (HFIP). The mixture was slowly dried to a thin film under nitrogen stream, followed by overnight lyophilization. The dried thin film was redissolved in 3 ml of 8 M urea containing ~20 mg 1,2-dihexanoyl-sn-Glycero-3-Phosphocholine ($DH_6PC$; protonated or deuterated from Avanti Polar Lipids), then dialyzed (MWCO 3.5 kDa) against 20 mM Tris buffer, pH 6.8 to remove the denaturant. During and after the dialysis, additional $DH_6PC$ was added to make up for the $DH_6PC$ loss due to dialysis, adjusting the DMPC:$DH_6PC$ ratio ($q$) to approximately 0.5. The bicelle $q$ was quantified by signal integration of the DMPC and $DH_6PC$ methyl peaks in the 1D $^1H$ NMR spectrum. The sample was concentrated in a Centricon (EMD Millipore, Billerica, MA) to ~350 μl. The final NMR sample contained ~0.8 mM MPER-TMD (monomer), ~55 mM DMPC, ~100 mM $DH_6PC$, 20 mM Tris (pH 6.8), 0.02% NaN3 and 5% $D_2O$.

Analysis of oligomeric state of the TMD of SARS-CoV-2 S2 in bicelles by SDS-PAGE

Wild type (WT) and mutants of $S2^{1209\text{-}1237}$ were first reconstituted in bicelles ($q$ = 0.5) and then mixed with SDS-PAGE sample loading buffer (Invitrogen) without boiling, followed by SDS-PAGE at 200 volts for 30 minutes and Commassie blue staining. Purified WT $S2^{1209\text{-}1237}$ (without reconstitution) migrated as monomer at ~5 kDa (theoretical MW = 3.4 kDa; see Fig. S1c). The WT $S2^{1209\text{-}1237}$ (after reconstitution) migrated at ~15 kDa (theoretical MW = 10.2 kDa; see Fig. S1c), consistent with the size of a trimer.

Analysis of oligomeric state of the TMD of SARS-CoV-2 S2 in bicelles by OG-label

In the OG-label method, each protomer of the oligomer to be studied is non-covalently labeled with a soluble cross-linkable protein (SCP), so that the latter can be cross-linked with Lomant's reagents to read out the sample oligomeric state. The small Ig-fold protein named GB1 (MW = 8.4 kDa) has been proven to serve as the SCP very effectively. A TriNTA molecule is linked via PEG-2-SMCC (succinimidyl 4-(N-maleimidomethyl)cyclohexane-1-carboxylate) to the N-terminus of GB1, while a $His_6$-tag is added to the C-terminus of the oligomeric protein, so that the TriNTA-GB1 conjugate can strongly attach to the protomer (the binding affinity of TriNTA to $His_6$-tag is 20 ± 10 nM). The GB1s are then cross-linked to report the oligomeric state of the protein, as the local concentration of stoichiometric amount of GB1 to the oligomer allows for more efficient cross-linking than for the free GB1 in solution. Finally, the cross-linked GB1s are released from the oligomer by addition of EDTA and analyzed by SDS-PAGE.

To implement the OG-label method for the $S2^{1209-1237}$, a $His_6$-tag was added at the C-terminus of the protein. The $His_6$-tagged protein was expressed, purified and reconstituted in bicelles ($q$ = 0.5) as described above, except the sample buffer was a phosphate buffer (pH 7.4) for better cross-linking efficiency. To prevent unwanted cross-linking between the membrane protein and GB1, all the free primary amines of the $S2^{1209-1237}$ were blocked by reacting with 100-fold molar excess of Sulfo-NHS acetate (Thermo Fisher Scientific) at room temperature for 1.5 hours. Excess Sulfo-NHS acetate was removed by dialysis while tightly controlling the bicelle $q$. After dialysis, the sample was concentrated to 30 μM and mixed with 60 μM TriNTA-GB1 to ensure that all the $His_6$-tags were saturated with TriNTA-GB1. The mixtures were then incubated at room temperature with various concentration of BS3(PEG9) (0.5, 1.0, and 2.0 mM, corresponding to lane 4, 5, and 6 in the gels shown in Fig. S1d) for 30 minutes, followed by a second incubation at room temperature with 0.6 mM glutaraldehyde for 3 minutes. The cross-linking reactions were quenched with 20 mM Tris buffer (pH 7.5) upon incubation at room temperature for 15 minutes. As negative control, 2.0 mM BS3(PEG9) and 0.6 mM glutaraldehyde were sequentially added to 60 μM TriNTA-GB1 in the absence of the $His_6$-tagged sample (lane 3 in the gels shown in Fig. S1d), indicating that the TriNTA-GB1 alone remains mostly monomeric in the conditions used. The cross-linked GB1s were then released from the samples by adding 50 mM EDTA and examined by SDS-PAGE using 12% Bis-Tris protein gels (Thermo Fisher Scientific) (Fig. S1d).

NMR resonances and NOE assignment

NMR data were collected at 303K on Bruker spectrometers operating at $^1H$ frequency of 900 MHz, 700 MHz, or 600 MHz equipped with cryogenic probes. NMR data sets were processed using nmrPipe[2]. NMR spectra were analyzed using Sparky (T. D. Goddard and D. G. Kneller, SPARKY 3, University of California, San Francisco) and XEASY[3]. Peak intensities were measured at peak local maxima using quadratic interpolation to identify peak centers. Origin (OriginLab, Northampton, MA) was used to fit the experimental data.

Sequence-specific assignment of backbone $^1H^N$, $^{15}N$, $^{13}C^{\alpha}$ and $^{13}C'$ resonances were accomplished using 3D TROSY-based HNCA, HN(CO)CA, HN(CA)CO and HNCO spectra[4-5], recorded using a ($^{15}N$, $^{13}C$, 90% $^2H$)-labeled sample. The aliphatic and aromatic resonances of the protein sidechains were assigned using a 3D $^{15}N$-edited NOESY-TROSY-HSQC ($\tau_{NOE}$ = 60 ms) and a 3D $^{13}C$-edited NOESY-HSQC ($\tau_{NOE}$ = 100 ms) spectra, recorded at $^1H$ frequency of

900 MHz. These NOESYs were performed using a ($^{15}N$, $^{13}C$)-labeled protein sample in bicelles ($q$ = 0.55) made of DMPC and $DH_6PC$ with deuterated acyl chains (Avanti Lipids). Specific stereo assignments of the methyl groups of valines and leucines were obtained from a $^1H$-$^{13}C$ HSQC spectrum recorded with 28 ms $^{13}C$ constant-time evolution, on a 700 MHz spectrometer, using a 15% $^{13}C$-labeled sample[6]. The same 3D $^{15}N$-edited NOESY-TROSY-HSQC and $^{13}C$-edited NOESY-HSQC spectra were used to assign local NOE-derived distance restraints.

For assigning inter-chain distance restraints, a $J_{CH}$-coupled NOE experiment was performed to exclusively detect inter-chain NOEs between the $^{15}N$-attached protons of one chain and the $^{13}C$-attached protons of the neighboring chains, using a mixed sample containing 50% ($^{15}N$, $^2H$)-labeled and 50% $^{13}C$-labeled protein. In this experiment, two interleaved 3D $^{15}N$-edited NOESY-TROSY spectra ($\tau_{NOE}$ = 150 ms) were recorded, at $^1H$ frequency of 900 MHz, in which one has $^{13}C$ decoupling during $^1H$ evolution before NOE mixing and the other does not.

Independent of the $J_{CH}$-coupled NOE experiment, we also performed the $J_{CH}$-modulated NOE experiment[1], in which the $^1H$ evolution period before the NOE mixing is changed to a "mixed constant-time" evolution to introduce $^1H$-$^{13}C$ $J$ evolution. Two interleaved spectra were recorded at $^1H$ frequency of 900 MHz with different $J_{CH}$ evolution ($J_{CH}$ = 0 ms, and $J_{CH}$ = 8 ms) before the NOE mixing ($\tau_{NOE}$ = 150 ms). For $J_{CH}$ = 0 ms, the inter- and intra- molecular NOE peaks are both positive, whereas for $J_{CH}$ = 8 ms, the inter- and intra- molecular NOE peaks are negative and positive, respectively. The $^1H$ transverse relaxation during 8 ms mixed constant-time evolution resulted in significant loss of sensitivity, and hence, a shorter S2 fragment with better relaxation property ($S2^{1215\text{-}1237}$) was used for this experiment.

NMR structure calculation

The structures were generated using the program XPLOR-NIH[7]. First, the monomer structure was generated using local NOE restraints and the backbone dihedral restraints derived from the backbone $^{15}N$, $^1H^N$, $^{13}C\alpha$, and $^{13}C'$ chemical shifts (using the TALOS+ program[8]). Second, the monomer structure and inter-chain NOE restraints were used with the ExSSO program[9] to generate a unique solution of trimeric assembly (Fig. S4a, S4b). Finally, the initial trimer solution was fed to the XPLOR-NIH for iterative refinement against all NMR restraints, while assigning more self-consistent inter-chain NOEs in both $^{13}C$-edited NOESY-HSQC and isotopically mixed NOE spectra after each iteration (Fig. S4c, S4d, S9).

For each inter-chain restraint between two adjacent chains, three identical distance restraints were assigned respectively to all pairs of neighboring chains to satisfy the condition of C3 rotational symmetry. The XPLOR refinement used a simulated annealing (SA) protocol in which the temperature in the bath was cooled from 1000 to 200 K with steps of 20 K. The NOE restraints were enforced by flat-well harmonic potentials, with the force constant ramped from 2 to 30 kcal/mol $Å^{-2}$ during annealing. Backbone dihedral angle restraints were taken from the 'GOOD' dihedral angles from TALOS+, all with a flat-well (±the corresponding uncertainties from TALOS+) harmonic potential with force constant ramped from 5 to 1000 kcal/mol $rad^{-2}$. A total of 75 structures were calculated and 15 lowest energy structures were selected as the final structural ensemble (Fig. S4e; Table S2).

Analysis of transmembrane partition

The paramagnetic probe titration (PPT) method[1, 10] was used to determine the membrane partition of the S2 TMD. This method is based on the notion that if the bicelle is sufficiently wide ($q$ > 0.5), the lateral solvent PRE becomes negligible, thus allowing the use of measurable solvent paramagnetic relaxation enhancement (PRE) to probe residue-specific depth immersion of the protein in the bilayer region of the bicelle. To ensure that our bicelles are wide enough, we reconstituted ($^{15}N$, $^{2}H$)-labeled S2$^{1209\text{-}1237}$ in bicelles with $q$ = 0.6 (Fig. S1b). The water-soluble and membrane-inaccessible paramagnetic agent, Gd-DOTA (Sigma), was used to generate solvent PRE. Gd-DOTA stock solution (200 mM) was titrated into the bicelle sample to reach final concentrations of 0.0, 1.0, 2.0, 4.0, 6.0, 8.0, 10.0 and 15.0 mM. At each concentration, a 2D $^{1}H$-$^{15}N$ TROSY-HSQC spectrum was recorded at 600 MHz to measure residue-specific PRE, defined here as the ratio of peak intensity in the presence (I) and absence ($I_0$) of the paramagnetic agent. Peak intensities were measured at peak local maxima using quadratic interpolation to identify peak centers. For each of the residues, we used *Origin* (OriginLab, Northampton, MA) to fit the PRE titration curve to exponential decay

$$\frac{I}{I_0} = 1 - PRE_{amp}\left(1 - e^{-\frac{[Gd-DOTA]}{\tau}}\right) \qquad \text{(Eq. S1)}$$

to derive the residue-specific PRE amplitude ($PRE_{\text{amp}}$) (Fig. S7a). To determine the position of the trimeric TMD relative to the bilayer center, we calculated, for each residue $i$, the distance ($r_Z$) along the protein symmetry axis from the amide proton to an arbitrary reference point based on the structure of the trimeric TMD. This calculation converted $PRE_{amp}$ vs. (residue number) to $PRE_{amp}$ vs. $r_Z$, which was then analyzed using the sigmoidal fitting method. Briefly, the trimer structure was moved along the 3-fold axis in increment of 0.5 Å (Fig. S8a, left) to achieve the best fit to the symmetric sigmoid equation:

$$PRE_{amp} = PRE_{amp}^{min} + \frac{\left(PRE_{amp}^{max} - PRE_{amp}^{min}\right)}{1+e^{\left(r_Z^I - |r_Z|\right)/SLOPE}} \qquad \text{(Eq. S2)}$$

where $PRE_{amp}^{min}$ and $PRE_{amp}^{max}$ are the limits within which $PRE_{amp}$ can vary for a particular protein system, $r_Z^I$ is the inflection point (the distance from the bilayer center at which $PRE_{amp}$ is halfway between $PRE_{amp}^{min}$ and $PRE_{amp}^{max}$), and *SLOPE* is a parameter which reports the steepness of the curve at the inflection point. The best fit (Fig. S8a, right) gave an adjusted coefficient of determination ($R^2_{adj}$) of 0.92 and was used to determine the position of the trimer structure with respect to the bilayer center ($r_Z$ = 0).

To complete the Gd-DOTA titration, a set of lipophilic PRE data was also acquired using a lipophilic paramagnetic agent 16-Doxyl-stearic acid (16-DSA). Using the identical approach to that employed for the solvent PRE, the bicelle-reconstituted S2$^{1209\text{-}1237}$ was titrated with the membrane-embedded paramagnetic agent 16-DSA at various known concentrations. The titrant stock solution (25 mM 16-DSA) was prepared in the same buffer as that of the protein sample, i.e., using $q$ = 0.6 bicelle solution to solubilize the paramagnetic agent to prevent changes in the sample bicelle $q$ during titration. The 16-DSA was added in small aliquots (8~10 μL per step) to minimize sample dilution. The progress of the titration was monitored by measuring a 2D $^{1}H$-$^{15}N$ TROSY-HSQC spectrum at each of the following 16-DSA concentrations: 0, 0.5, 1.0, 2.0, 3.0, 4.0, 5.0 and 7.5 mM. The residue-specific $PRE_{amp}$ was determined by fitting the peak intensity decay as a function of [16-DSA] to the exponential decay equation Eq. S1. The $PRE_{\mathrm{amp}}$ vs. (residue number) plot for 16-DSA shows a reciprocal profile of that of Gd-DOTA (Fig. S7b).

Hydrogen-deuterium (H-D) exchange

For measuring H-D exchange of the S2 TMD in bicelles, $S2^{1209\text{-}1237}$ was reconstituted in protonated solvent in the beginning and was flash-frozen in liquid nitrogen and then thoroughly lyophilized to get rid of protonated water. The dried sample was dissolved in 360 µL of 99.9% $D_2O$. The progress of the H-D exchange was monitored by measuring a 2D $^1$H-$^{15}$N TROSY-HSQC spectrum at uniform time intervals of ~3 hours up to ~5 days. The residue-specific exchange constant, $k_{ex}$ (=1/$\tau_{ex}$), was determined by fitting the fractional peak intensity vs. time to the following exponential decay equation:

$$\frac{I}{I_0} \propto \left(e^{-\frac{t}{\tau ex}}\right) \qquad \text{(Eq. S3)}$$

where $I_0$ and $I$ are the peak intensities before and after the H-D exchange, $t$ is the time passed from the beginning of the exchange, and $\tau_{ex}$ is the time constant of the decay.

NMR Dynamics Measurements

Residue-specific rotational correlation time ($\tau_C$) was measured using a 2D version of the 1D experiment known as TRACT (TROSY for rotational correlation times)[11]. For this experiment, a sample of ~0.9 mM ($^{15}$N, $^{13}$C, 90% $^2$H)-labeled $S2^{1209\text{-}1237}$ reconstituted in bicelles with $q$ = 0.55 was used. In the TRACT experiment, $^{15}$N relaxation delays were set to 0, 8, 16, 32, 40, 56, 80 ms for the TROSY component, and set to 0, 8, 16, 20, 24, 32, 50 ms for anti-TROSY component. Due to the fast relaxation of the anti-TROSY component, 4-fold more scans were used for recording the anti-TROSY spectra. For each of the assigned peaks, the $R_2$ relaxation rate of the TROSY ($R_2^{\alpha}$) and anti-TROSY ($R_2^{\beta}$) components were determined by fitting the measurements to the exponential decay function:

$$I/I_0 = y + Ae^{-R*Delay} \qquad \text{(Eq. S4)}$$

where $I$ and $I_0$ represent the peak intensity with and without relaxation delay, respectively. Data fitting was done using the Origin program (OriginLab, Northampton, MA). $\tau_C$ was derived from the following equations:

$$R_\beta - R_\alpha = 2p\delta_N\big(4\,J(0) + 3\,J(\omega_N)\big)(3cos^2\theta - 1) \qquad \text{(Eq. S5)}$$

$$J(\omega) = 0.4\tau_C/[1 + (\tau_C\omega)^2] \qquad \text{(Eq. S6)}$$

where $p$ is the dipole-dipole coupling between $^1$H and $^{15}$N and $\delta_N$ is the chemical shift anisotropy of the $^{15}$N nucleus with the parameter $\theta$ = 17°, and ω is the spectrometer $^{15}$N frequency (60 MHz).

### Table S1. Reported structures of SARS-CoV-2 spike protein in Year 2020

| PDB ID | Method | Construct | Structured region | state | Reference |
|---|---|---|---|---|---|
| 6LXT | X-ray | 910-1206 | 913-988, 1163-1202 | Post-fusion state | 12 |
| 6VSB | EM | 1-1208 | 27-1146 | Ecto-domain trimer in pre-fusion state | 13 |
| 6VXX | EM | 14-1211 | 27-1147 | Ecto-domain trimer in close state | 14 |
| 6VYB | EM | 14-1211 | 27-1147 | Ecto-domain trimer in open state | 14 |
| 6X29 | EM | 16-1208 | 27-1147 | Ecto-domain trimer in rS2d Down State | 15 |
| 6X2A | EM | 16-1208 | 27-1147 | Ecto-domain trimer in u1S2q 1 Up State | 15 |
| 6X2B | EM | 16-1208 | 27-1147 | U1S2q 2 RBD Up Spike Protein Trimer | 15 |
| 6X2C | EM | 16-1208 | 27-1147 | U1S2q All Down RBD State | 15 |
| 6X79 | EM | 14-1211 | 27-1140 | Ecto-domain trimer in close state | 16 |
| 6XR8 | EM | 1-1273 | 14-1162 | S protein trimer in prefusion state | 17 |
| 6XRA | EM | 1-1273 | 703-1197 | S protein trimer in postfusion state | 17 |
| 6XS6 | EM | 1-1213 | 27-1147 | Spike D614G variant | 18 |
| 6ZGE | EM | 1-1208 | 27-1146 | S protein in closed state | 19 |
| 6ZGF | EM | 1-1208 | 27-1146 | S protein in closed state | 19 |
| 6ZGG | EM | 1-1208 | 27-1146 | Furin Cleaved Spike Protein | 19 |
| 6ZGH | EM | 1-1208 | 27-1146 | Intermediate state | 19 |
| 6ZGI | EM | 1-1208 | 27-1146 | Closed state | 19 |
| 6ZOX | EM | 14-1211 | 27-1146 | Disulphide-stabilized Spike Protein Trimer | 20 |
| 6ZOY | EM | 14-1211 | 27-1146 | Disulphide-stabilized Spike Protein Trimer | 20 |
| 6ZOZ | EM | 14-1211 | 27-1146 | Disulphide-stabilized Spike Protein Trimer | 20 |
| 6ZP0 | EM | 14-1211 | 27-1146 | Disulphide-stabilized Spike Protein Trimer | 20 |
| 6ZP1 | EM | 14-1211 | 27-1146 | Disulphide-stabilized Spike Protein Trimer | 20 |
| 6ZP2 | EM | 14-1211 | 27-1146 | Disulphide-stabilized Spike Protein Trimer | 20 |
| 6ZWV | EM | 1-1273 | 27-1151 | Spike Proteins on intact virions | 21 |
| 7JJI | EM | 1-1273 | 14-1146 | Prefusion spike trimer | 22 |
| 7JJJ | EM | 1-1273 | 14-1146 | Dimers of spike trimers | 22 |

**Table S2. NMR and refinement statistics for the TMD of SARS-CoV-2 S protein**

| **NMR distance and dihedral constraints** [a] | TMD (1217-1237) |
|---|---|
| Distance constraints from NOE | 387 |
| Short-range intramolecular ($\|i-j\| \leq 4$) | 109 x 3 |
| Long-range intramolecular ($\|i-j\| \geq 5$) | 0 |
| Intermolecular | 20 x 3 |
| Total dihedral angle restraints [b] | 120 |
| $\phi$ (TALOS) | 20 x 3 |
| $\psi$ (TALOS) | 20 x 3 |
| **Structure statistics** [c] | |
| Violations (mean ± s.d.) | |
| Distance constraints (Å) | 0.091 ± 0.005 |
| Dihedral angle constraints (°) | 0.210 ± 0.042 |
| Deviations from idealized geometry | |
| Bond lengths (Å) | 0.006 ± 0.000 |
| Bond angles (°) | 0.683 ± 0.020 |
| Impropers (°) | 0.367 ± 0.020 |
| Average pairwise r.m.s. deviation (Å) [d] | |
| Heavy | 0.958 |
| Backbone | 0.579 |

[a] The numbers of constraints are summed over all three subunits.

[b] Backbone $\phi$ and $\psi$ restraints and their respective uncertainties were obtained from the "GOOD" dihedrals generated by the TALOS+ program[8] based on the backbone chemical shift values.

[c] Statistics are calculated and averaged over an ensemble of the 15 lowest energy structures out of 75 calculated structures.

[d] The precision of the atomic coordinates is defined as the average r.m.s. difference between the 15 final structures and their mean coordinates. The atomic structure coordinate and structural constraints have been deposited in the Protein Data Bank (PDB), accession number 7LC8. The chemical shift values have been deposited in the Biological Magnetic Resonance Data Bank (BMRB), accession number 30842.

**Table S3. Residue specific $PRE_{amp}$ of S2 TMD in $q$ = 0.6 bicelles**

| Residue | $PRE_{amp}$ (Gd-DOTA) | $PRE_{amp}$ (16-DSA) |
|---|---|---|
| 1218 | 0.67±0.028 | 0.79±0.047 |
| 1219 | 0.61±0.034 | 0.84±0.044 |
| 1220 | 0.53±0.045 | 0.83±0.052 |
| 1221 | 0.56±0.036 | 0.84±0.051 |
| 1222 | 0.57±0.042 | 0.87±0.048 |
| 1223 | 0.58±0.037 | 0.84±0.036 |
| 1224 | 0.53±0.029 | 0.85±0.026 |
| 1225 | 0.59±0.044 | 0.88±0.042 |
| 1226 | 0.52±0.047 | 0.86±0.044 |
| 1227 | 0.54±0.038 | 0.89±0.043 |
| 1228 | 0.60±0.021 | 0.85±0.046 |
| 1229 | 0.57±0.027 | 0.82±0.049 |
| 1230 | 0.58±0.035 | 0.87±0.045 |
| 1231 | 0.60±0.042 | 0.82±0.044 |
| 1232 | 0.56±0.051 | 0.85±0.032 |
| 1233 | 0.60±0.046 | 0.84±0.055 |
| 1234 | 0.71±0.043 | 0.81±0.053 |
| 1235 | 0.73±0.056 | 0.78±0.052 |
| 1236 | 0.82±0.052 | 0.68±0.058 |
| 1237 | 0.89±0.067 | 0.58±0.062 |

**Table S4. Membrane localization of S2 TMD trimer by PPT analysis**

| Residue ($H^N$) | $r_z$ (Å) |
|---|---|
| L1218 | -11.63689 |
| G1219 | -9.42969 |
| F1220 | -8.89779 |
| I1221 | -8.22165 |
| A1222 | -6.05935 |
| G1223 | -4.217 |
| L1224 | -3.97906 |
| I1225 | -3 |
| A1226 | -0.73905 |
| (Membrane center) | 0.0 |
| I1227 | 0.90939 |
| V1228 | 1.53238 |
| L1229 | 2.89463 |
| V1230 | 5.13462 |
| T1231 | 6.3745 |
| I1232 | 7.21047 |
| L1233 | 9.06871 |
| L1234 | 10.98703 |
| S1235 | 11.6717 |
| S1236 | 12.88747 |
| T1237 | 15.15529 |

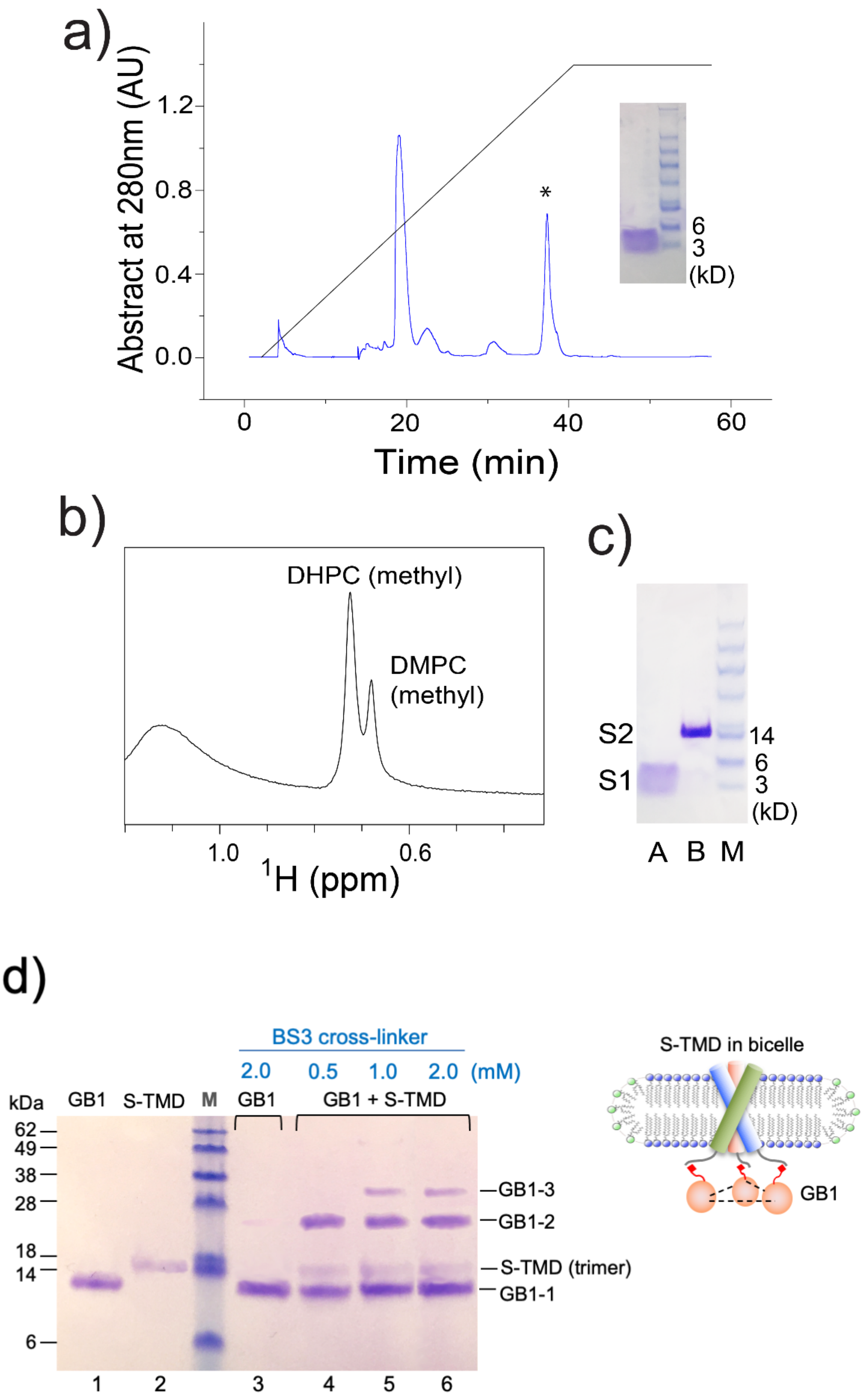
a)
Abstract at 280nm (AU)
1.2
0.8
0.4
0.0
0
20
40
60
Time (min)
*
6
3
(kD)
b)
DHPC (methyl)
DMPC
(methyl)
1.0
0.6
1H (ppm)
c)
S2
S1
14
6
3
(kD)
A B M
d)
BS3 cross-linker
2.0 0.5 1.0 2.0 (mM)
kDa
GB1
S-TMD
M
GB1
GB1 + S-TMD
62
49
38
28
18
14
6
GB1-3
GB1-2
S-TMD (trimer)
GB1-1
1 2 3 4 5 6
S-TMD in bicelle
GB1

**Figure S1. Purification and bicelle reconstitution of SARS-CoV-2 $S2^{1209-1237}$**

(**a**) Reverse phase HPLC purification of $S2^{1209-1237}$ from CNBr-cleaved trpLE-$S2^{1209-1237}$ fusion protein on Zorbax SB-C3 column with a gradient from 95% $dH_2O$, 5% isopropanol (IPA), 0.1% trifluoroacetic acid (TFA) (buffer A) to 75% IPA, 25% acetonitrile, 0.1% TFA (buffer B). The product was verified by SDS-PAGE and Mass Spectrometry.

(**b**) $^1H$ NMR spectrum of the reconstituted bicelle sample recorded at 600 MHz, showing the molar ratio of DMPC to $DH_6PC$ to be ~0.55.

(**c**) Oligomerization of $S2^{1209-1237}$ in bicelles analyzed by SDS-PAGE. The gel lanes from left to right are: A – purified $S2^{1209-1237}$ powder without reconstitution; B – $S2^{1209-1237}$ reconstituted in DMPC-$DH_6PC$ bicelles ($q$ = 0.55); M – MW markers. Both samples were dissolved in gel loading buffer prior to SDS-PAGE. S1: Monomer band at ~5 kDa (theoretical MW = 3.4 kDa). S2: Trimer band at ~15 kDa (theoretical MW = 10.2 kDa).

(**d**) Oligomerization of $S2^{1209-1237}$ in bicelles analyzed by the OG-label method[23] that uses the GB1 protein as reporter of $S2^{1209-1237}$ oligomerization. The SDS-PAGE gel lanes are: 1 – GB1 (GB1 conjugated to a triNTA molecule); 2 – $S2^{1209-1237}$ in DMPC-$DH_6PC$ bicelles with $q$ = 0.55 (trimeric in SDS-PAGE); 3 – GB1 alone cross-linked with 2 mM BS3 cross-linker; 4-6 – GB1 cross-linked in the presence of bicelle-reconstituted $S2^{1209-1237}$ at increasing cross-linker concentration.

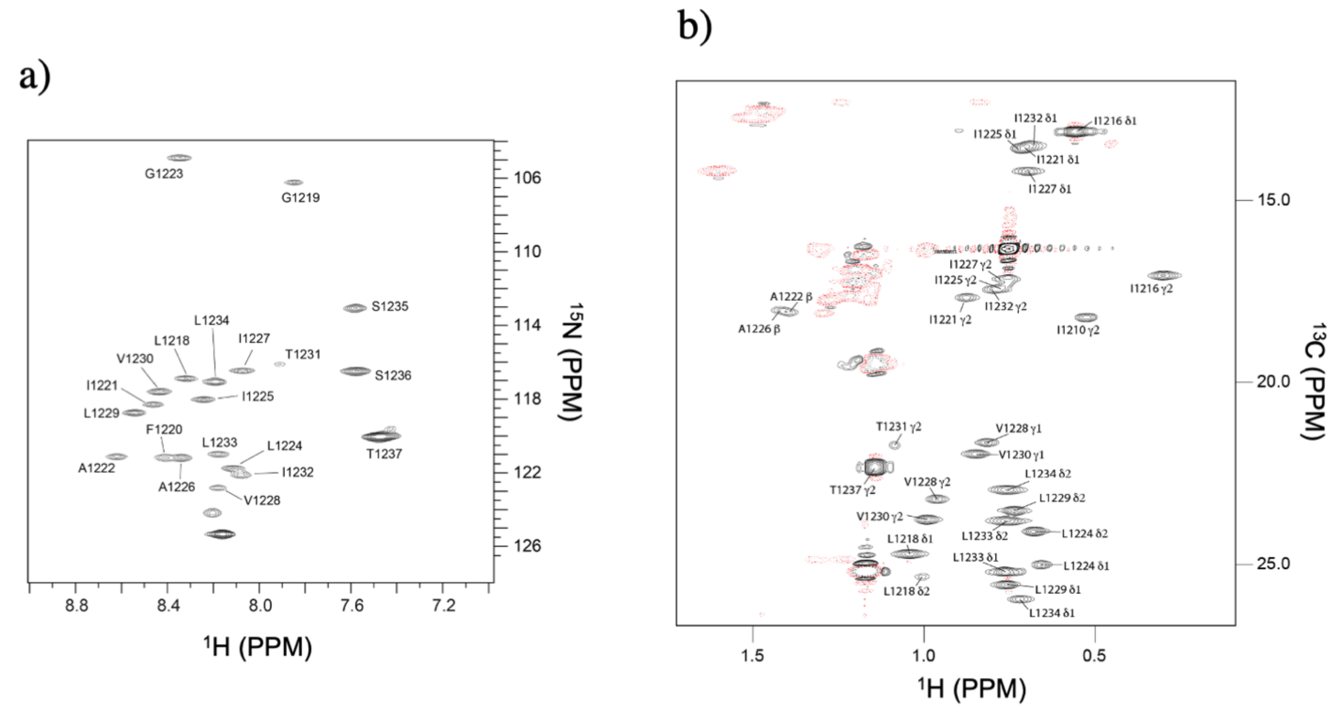


**Figure S2. NMR spectra of S2$^{1209\text{-}1237}$ reconstituted in DMPC-DH$_6$PC bicelles with *q* = 0.55**

(**a**) The $^1$H-$^{15}$N TROSY-HSQC spectrum of ($^{15}$N,$^{13}$C, $^2$H)-labeled S2$^{1209\text{-}1237}$, recorded at $^1$H frequency of 600 MHz at 303 K.

(**b**) The methyl group region of the 2D $^1$H-$^{13}$C HSQC spectrum (28 ms constant-time $^{13}$C evolution), recorded at $^1$H frequency of 700 MHz using ($^{15}$N, $^{13}$C)-labeled S2$^{1209\text{-}1237}$ reconstituted in DMPC and DH$_6$PC with deuterated acyl chains.

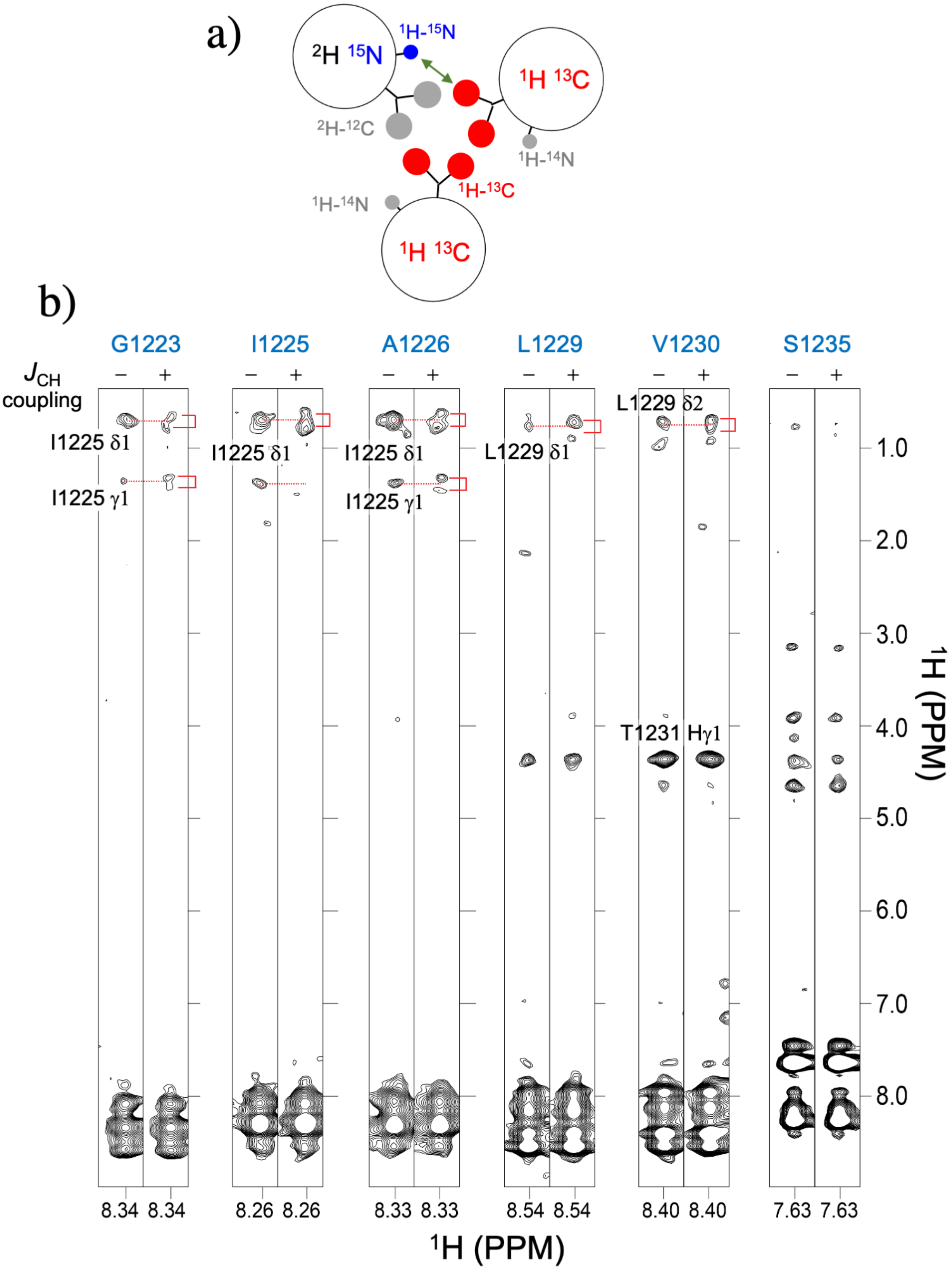
a)
2H 15N
1H-15N
1H 13C
2H-12C
1H-14N
1H-13C
1H-14N
1H 13C
b)
G1223
I1225
A1226
L1229
V1230
S1235
JCH coupling
− +
I1225 δ1
I1225 γ1
I1225 δ1
I1225 δ1
I1225 γ1
L1229 δ1
L1229 δ2
T1231 Hγ1
8.34 8.34
8.26 8.26
8.33 8.33
8.54 8.54
8.40 8.40
7.63 7.63
1H (PPM)
1.0
2.0
3.0
4.0
5.0
6.0
7.0
8.0
1H (PPM)

c)

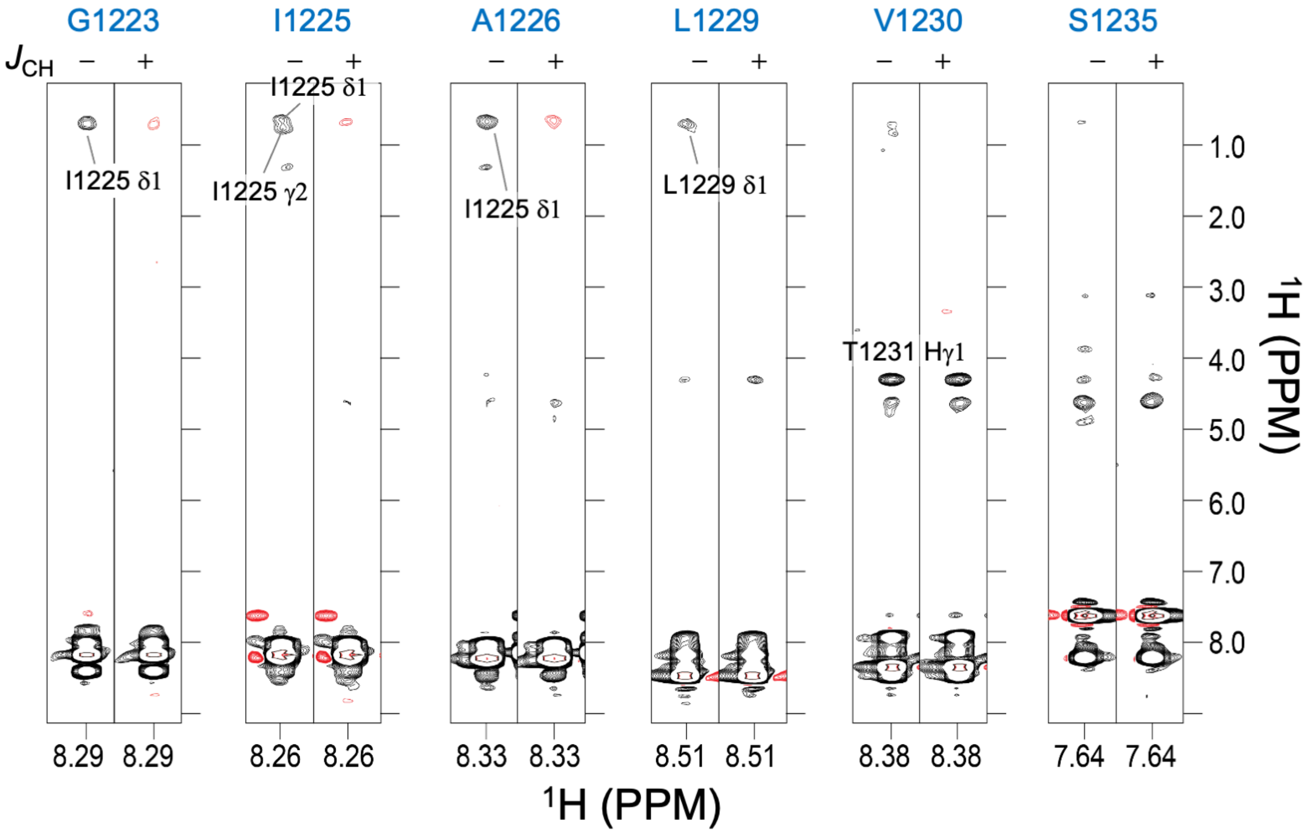


**Figure S3**. **Detection of inter-chain NOEs**

(**a**) Schematic illustration of the mixed sample comprising 1:1 ratio of ($^{15}$N, $^{2}$H)-labeled S2$^{1209-1237}$ to $^{13}$C-labeled S2$^{1209-1237}$. $^{1}$H-$^{15}$N and $^{1}$H-$^{13}$C groups are shown in blue and red, respectively, whereas the undetectable groups (e.g., $^{1}$H-$^{14}$N and $^{2}$H-$^{12}$C) are shown in gray.

(**b**) Residue-specific stripes from a 3D $^{15}$N-edited NOESY-TROSY ($\tau_{NOE}$ = 150 ms), recorded in an interleaved fashion with and without $^{13}$C decoupling during $^{1}$H evolution before NOE mixing. The spectrum was recorded at 900 MHz and 303 K using the mixed sample in (a). For each of the selected residues, two stripes are shown: left – $^{13}$C decoupled; right – $^{13}$C coupled. The red brackets indicate peak splitting due to $J_{CH}$.

(**c**) Independent validation of inter-chain NOEs in (b) by the $J_{CH}$-modulated NOESY[1]. Residue-specific stripes from the 3D $J_{CH}$-modulated NOESY (NOE mixing time = 150 ms) recorded at 900 MHz and 303 K. For each of the selected residues, two stripes are shown: left – $J_{CH}$ = 0, inter- and intra- chain NOE peaks are both positive (black); right – $J_{CH}$ = 8 ms, inter-chain NOE peaks are negative (red). Note that the negative NOE peaks are significantly weaker due to cross correlation effects resulting from C-H dipolar interaction.

a

| Amide | Aliphatic | dist (Å) |
|---|---|---|
| I1221 HN | I1221 Hδ1 | 4.0 ± 1 |
| A1222 HN | I1221 Hδ1 | 4.0 ± 1 |
| G1223 HN | I1225 Hδ1 | 4.0 ± 1 |
| G1223 HN | I1225 Hγ2 | 4.0 ± 1 |
| I1225 HN | I1225 Hδ1 | 4.0 ± 1 |
| I1225 HN | I1225 Hγ2 | 4.0 ± 1 |
| A1226 HN | I1225 Hδ1 | 4.0 ± 1 |
| V1230 HN | L1229 Hδ2 | 4.0 ± 1 |

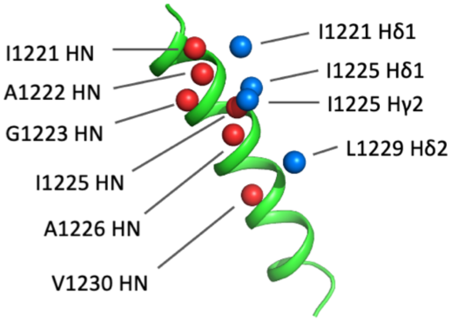


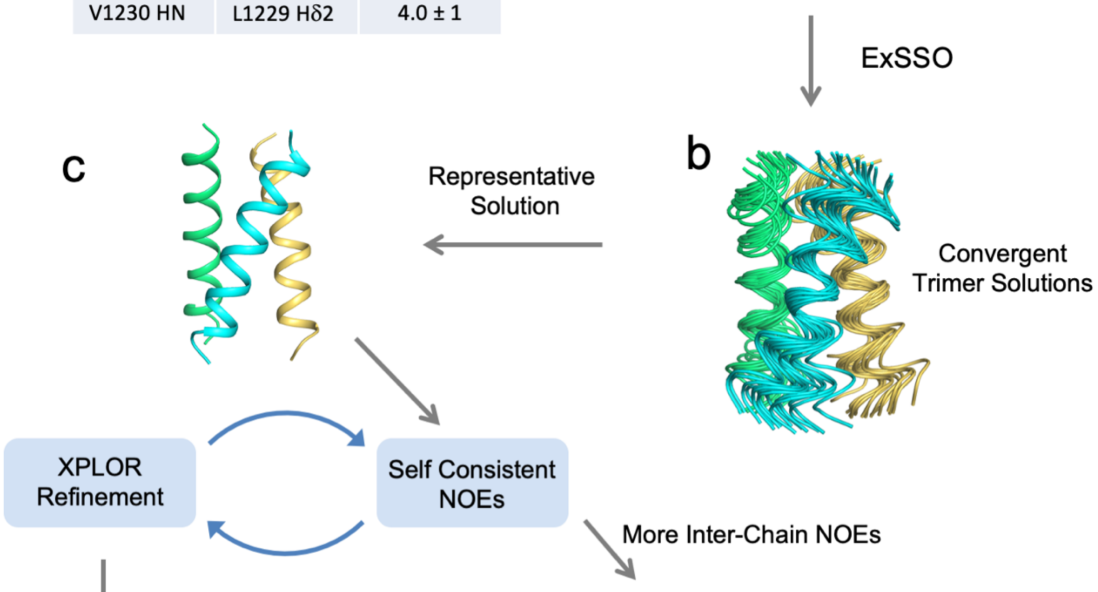


d

| Proton 1 | Proton 2 | Protomer | dist (Å) |
|---|---|---|---|
| L1218 Hδ1 | I1221 Hδ1 | i, i - 1 | 2.5 ± 1 |
| L1218 Hδ1 | F1220 Hε | i, i - 1 | 3.0 ± 1 |
| I1221 Hδ1 | L1218 Hδ1 | i, i + 1 | 2.5 ± 1 |
| I1225 Hδ1 | A1222 Hβ | i, i + 1 | 3.0 ± 1 |
| I1225 Hγ2 | A1226 Hα | i, i + 1 | 2.5 ± 1 |
| I1227 HN | L1229 Hδ2 | i, i - 1 | 4.0 ± 1 |
| V1228 Hγ1 | A1226 Hβ | i, i + 1 | 3.0 ± 1 |
| L1229 Hδ1 | A1226 Hβ | i, i + 1 | 2.5 ± 1 |
| L1229 Hδ2 | A1226 Hβ | i, i + 1 | 2.5 ± 1 |
| L1229 Hδ1 | I1227 HN | i, i + 1 | 4.0 ± 1 |
| L1229 Hδ2 | I1227 Hα | i, i + 1 | 4.0 ± 1 |
| V1230 HN | I1232 Hγ1 | i, i - 1 | 4.0 ± 1 |

e

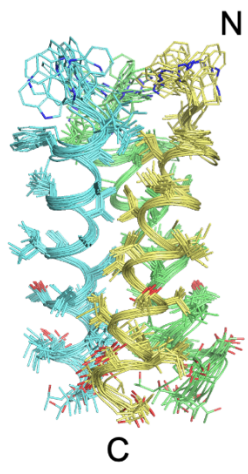


**Figure S4. Overall procedure of structure determination by NMR.**

(**a**) *Left*: Initial set of inter-chain NOE restraints identified in the NOE spectra of mixed isotope labeled sample (Fig. S3). *Right*: Protons associated with these NOEs are indicated by spheres in the context of the monomer structure generated with local NOE restraints and backbone dihedral restraints from TALOS+[8].

(**b**) The monomer structure and inter-chain NOE restraints (with ambiguous directionality) were subject to an exhaustive conformational search algorithm (ExSSO[9]), which found an ensemble of convergent solutions of trimeric assembly.

(**c**) A representative trimer structure from (b) was fed to the XPLOR-NIH[7] for iterative refinement against all NMR restraints, while assigning more self-consistent NOEs after each iteration.

(**d**) Additional list of inter-chain NOEs assigned during the iterative refinement step in (c) (also see Fig. S9 below).

(**e**) Ensemble of 15 lowest energy structures from 75 structures calculated in the final refinement step in (c). Protons are not displayed. The region displayed include residues 1217-1237. Residues 1209-1216 is unstructured in our sample and essentially no NMR data was collected for this region.

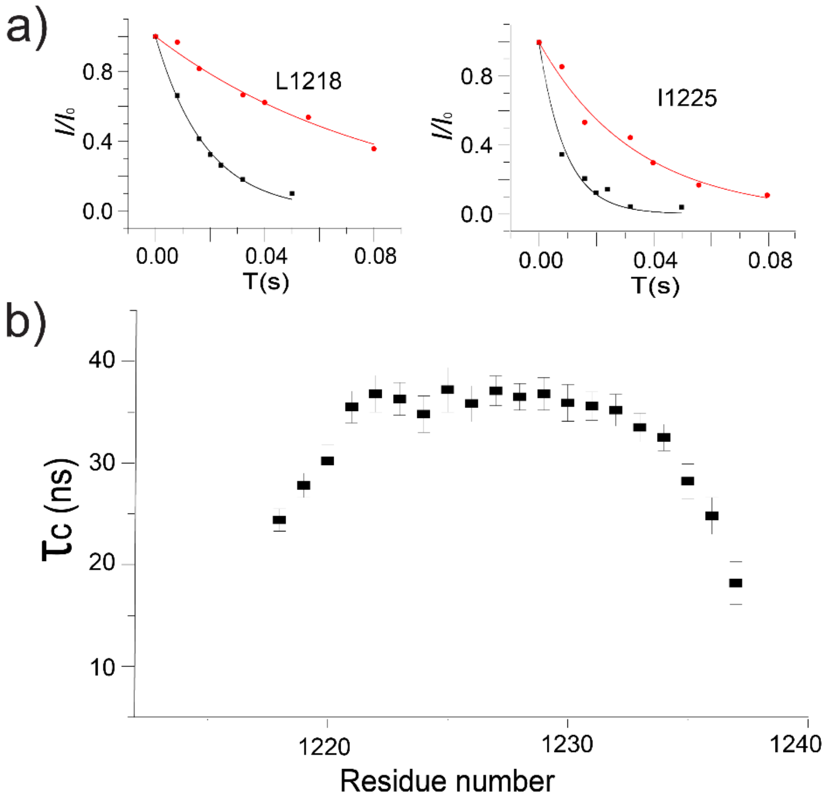


**Figure S5. Measurement of residue-specific $\tau_C$ for the S2 TMD in bicelles (*q* = 0.55) using the TRACT experiment**. The TRACT experiment[11] was performed in 2D $^1$H-$^{15}$N correlation mode to resolve resonance overlap. The data was collected at 600 MHz and 303K.

(**a**) Example plots showing decay of relative peak intensity, $I/I_0$, due to $^{15}$N transverse relaxation for the TROSY (red) and anti-TROSY (black) components. The TROSY relaxation rate ($R_\alpha$) and the anti-TROSY relaxation rate ($R_\beta$) were obtained by exponential fitting. $\tau_C$ was calculated using the difference $R_\beta$ - $R_\alpha$ as described in Ref [11].

(**b**) Residues-specific $\tau_C$ calculated for the S2 TMD in bicelles.

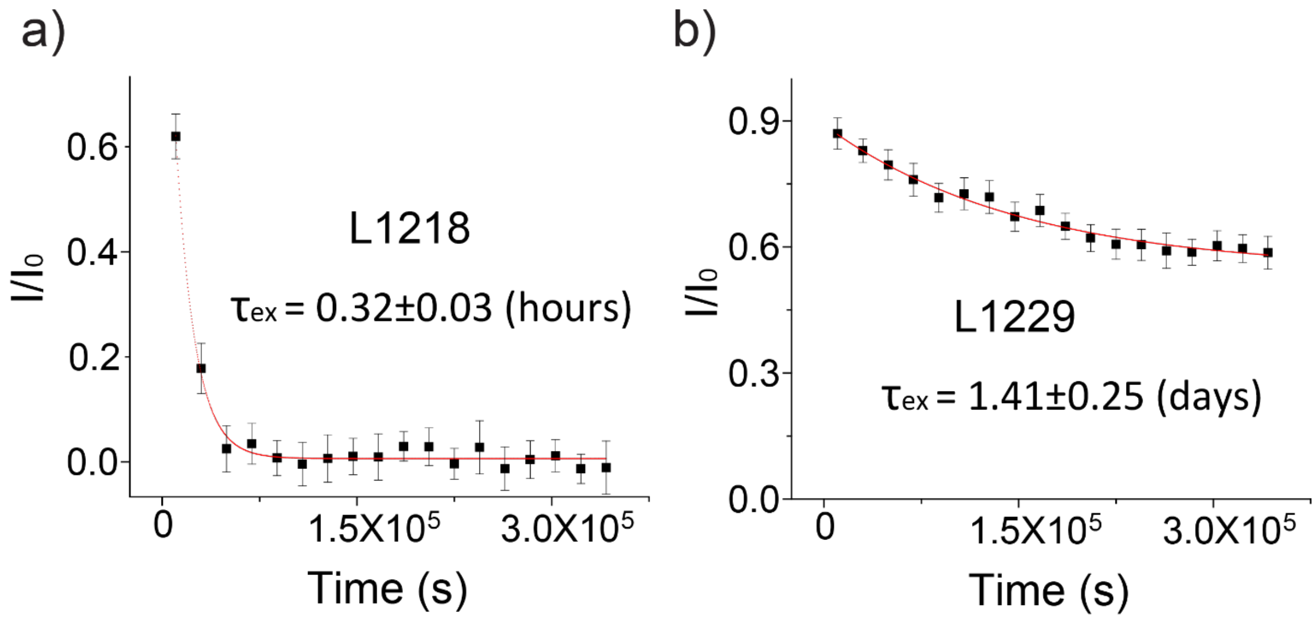


**Figure S6. H-D exchange of S2$^{1209\text{-}1237}$ in bicelles with *q* = 0.55**.

(**a**) Signal decay over time for L1218, representing the high exchange regime (the N-terminal end of the TM helix with higher solvent accessibility).

(**b**) Signal decay over time for L1229, representing the low exchange regime (the hydrophobic core).

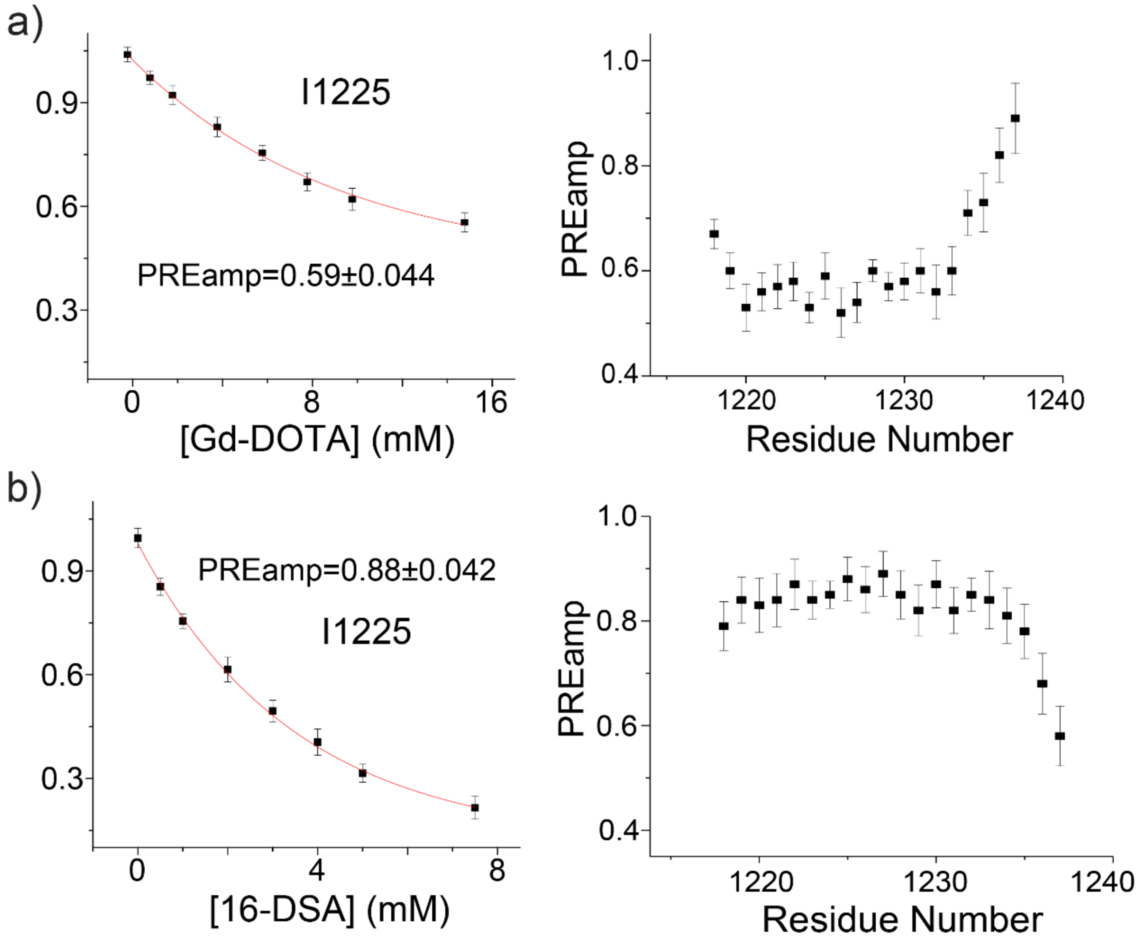


**Figure S7. Membrane partition analysis of the S2 TMD trimer**

(**a**) Solvent PRE titration with Gd-DOTA. $^{1}H$-$^{15}N$ TROSY-HSQC signal decay vs. Gd-DOTA concentration were fitted to Eq. S1 to determine the residue-specific $PRE_{amp}$ as shown for residue I1225 (left) and all residues (right).

(**b**) Lipophilic PRE titration with 16-DSA was performed to provide a reciprocal profile of $PRE_{amp}$ vs. (residue number), as shown for I1225 (left) and for all residues (right).

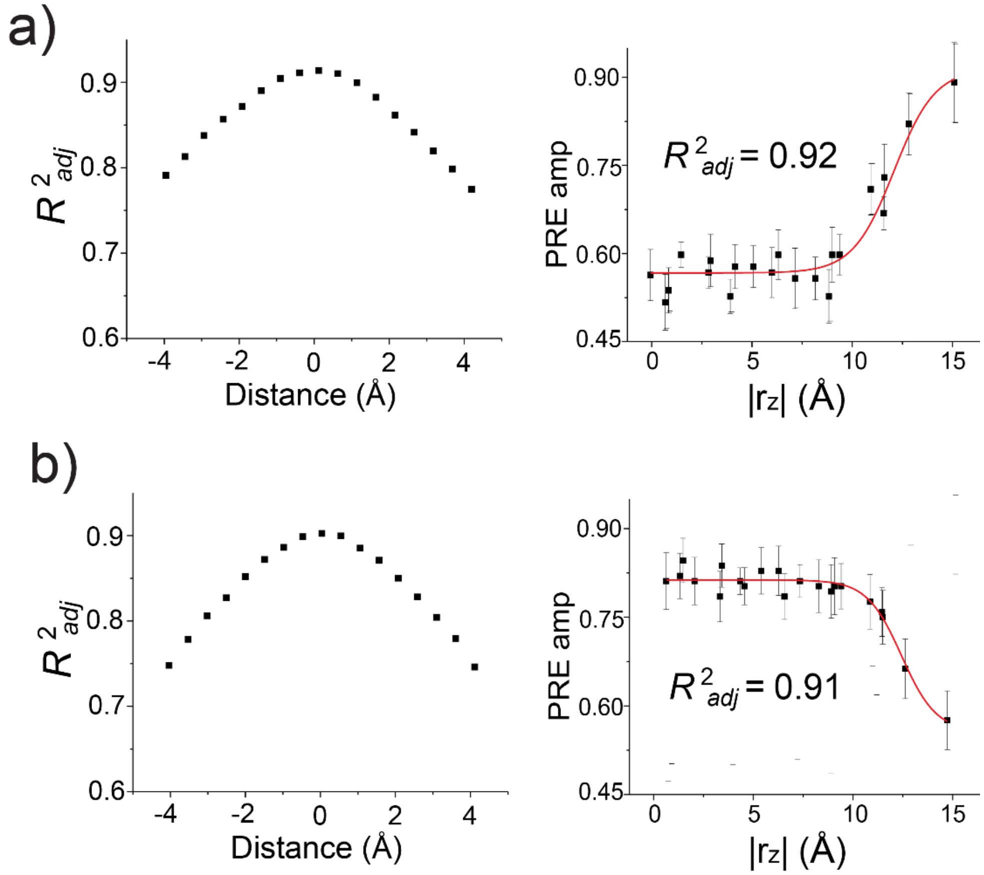


**Figure S8. Assignment of the bilayer center to the S2 TMD trimer using the paramagnetic probe titration (PPT) method**

(**a**) *Left*: Sliding of the S2 TMD trimer structure along the 3-fold axis (or bilayer normal) to yield best fit of Gd-DOTA generated $PRE_{amp}$ to the symmetric sigmoidal function (Eq. S2). *Right*: The $PRE_{amp}$ vs. $r_Z$ profile from the best fit, showing the adjusted coefficient of determination ($R^2_{adj}$).

(**b**) Same as in (a), but for the 16-DSA data set.

The plots show that $R^2_{adj}$ is a reliable indicator of the protein position with an error of about ± 0.5 Å. Both fittings show very close bilayer center position, which corresponds approximately to A1226/I1227.

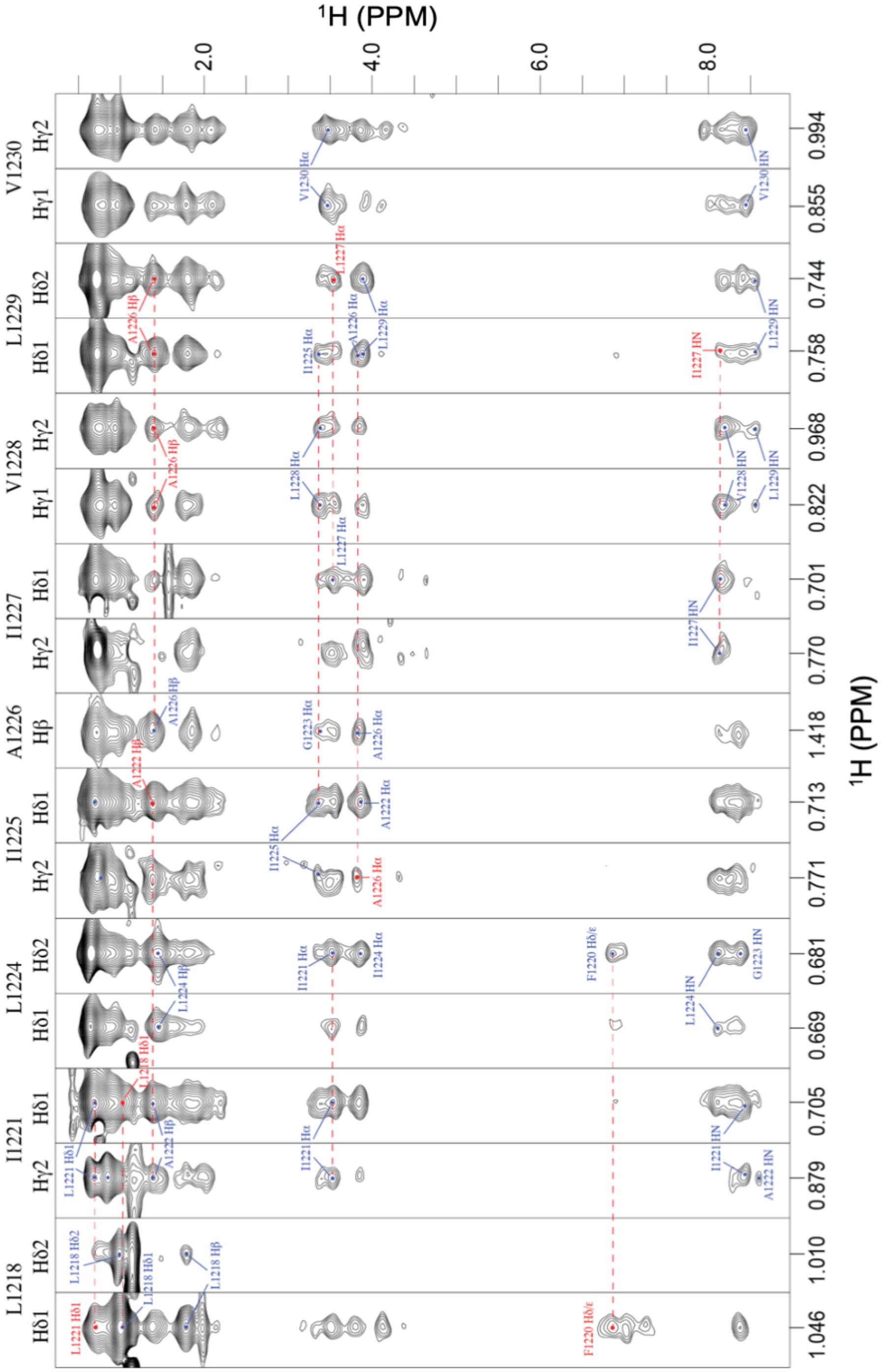


**Figure S9.** Additional inter-chain NOEs consistent with the trimer solution in Fig. S4b. Example stripe from 3D $^{13}$C-edited NOESY-HSQC spectrum recorded at 700 MHz ($^1$H frequency) and 303 K with a NOE mixing time of 100 ms. The red dashed lines indicate matching chemical shift between intra-monomer (blue) and inter-monomer (red) NOEs. The spectrum was acquired using ($^{15}$N,$^{13}$C)-labeled S2$^{1209\text{-}1237}$ reconstituted in bicelles with deuterated DMPC and DH$_6$PC acyl chains.

1. Fu, Q.; Piai, A.; Chen, W.; Xia, K.; Chou, J. J., Structure determination protocol for transmembrane domain oligomers. *Nat Protoc* **2019,** *14* (8), 2483-2520.
2. Delaglio, F.; Grzesiek, S.; Vuister, G. W.; Zhu, G.; Pfeifer, J.; Bax, A., NMRPipe: A multidimensional spectral processing system based on UNIX pipes. *Journal of Biomolecular NMR* **1995,** *6* (3), 277-293.
3. Bartels, C.; Xia, T. H.; Billeter, M.; Guntert, P.; Wuthrich, K., The program XEASY for computer-supported NMR spectral analysis of biological macromolecules. *J Biomol NMR* **1995,** *6* (1), 1-10.
4. Kay, L. E.; Ikura, M.; Tschudin, R.; Bax, A., Three-dimensional triple-resonance NMR Spectroscopy of isotopically enriched proteins. 1990. *J Magn Reson* **1990,** *213* (2), 423-41.
5. Salzmann, M.; Wider, G.; Pervushin, K.; Senn, H.; Wuthrich, K., TROSY-type triple-resonance experiments for sequential NMR assignments of large proteins. *Journal of the American Chemical Society* **1999,** *121* (4), 844-848.
6. Szyperski, T.; Neri, D.; Leiting, B.; Otting, G.; Wuthrich, K., Support of 1H NMR assignments in proteins by biosynthetically directed fractional 13C-labeling. *J. Biomol. NMR* **1992,** *2* (4), 323-334.
7. Schwieters, C. D.; Kuszewski, J.; Tjandra, N.; Clore, G. M., The Xplor-NIH NMR molecular structure determination package. *J. Magn. Reson.* **2002,** *160*, 66-74.
8. Shen, Y.; Delaglio, F.; Cornilescu, G.; Bax, A., TALOS+: a hybrid method for predicting protein backbone torsion angles from NMR chemical shifts. *Journal of Biomolecular NMR* **2009,** *44* (4), 213-23.
9. Yang, J.; Piai, A.; Shen, H. B.; Chou, J. J., An Exhaustive Search Algorithm to Aid NMR-Based Structure Determination of Rotationally Symmetric Transmembrane Oligomers. *Sci Rep* **2017,** *7* (1), 17373.
10. Piai, A.; Fu, Q.; Dev, J.; Chou, J. J., Optimal Bicelle Size q for Solution NMR Studies of the Protein Transmembrane Partition. *Chemistry* **2017,** *23* (6), 1361-1367.
11. Lee, D.; Hilty, C.; Wider, G.; Wuthrich, K., Effective rotational correlation times of proteins from NMR relaxation interference. *J Magn Reson* **2006,** *178* (1), 72-6.
12. Xia, S.; Liu, M.; Wang, C.; Xu, W.; Lan, Q.; Feng, S.; Qi, F.; Bao, L.; Du, L.; Liu, S.; Qin, C.; Sun, F.; Shi, Z.; Zhu, Y.; Jiang, S.; Lu, L., Inhibition of SARS-CoV-2 (previously 2019-nCoV) infection by a highly potent pan-coronavirus fusion inhibitor targeting its spike protein that harbors a high capacity to mediate membrane fusion. *Cell Res* **2020,** *30* (4), 343-355.
13. Wrapp, D.; Wang, N.; Corbett, K. S.; Goldsmith, J. A.; Hsieh, C. L.; Abiona, O.; Graham, B. S.; McLellan, J. S., Cryo-EM structure of the 2019-nCoV spike in the prefusion conformation. *Science* **2020,** *367* (6483), 1260-1263.
14. Walls, A. C.; Park, Y. J.; Tortorici, M. A.; Wall, A.; McGuire, A. T.; Veesler, D., Structure, Function, and Antigenicity of the SARS-CoV-2 Spike Glycoprotein. *Cell* **2020,** *181* (2), 281-292 e6.
15. Henderson, R.; Edwards, R. J.; Mansouri, K.; Janowska, K.; Stalls, V.; Gobeil, S. M. C.; Kopp, M.; Li, D.; Parks, R.; Hsu, A. L.; Borgnia, M. J.; Haynes, B. F.; Acharya, P., Controlling the SARS-CoV-2 spike glycoprotein conformation. *Nat Struct Mol Biol* **2020,** *27* (10), 925-933.
16. McCallum, M.; Walls, A. C.; Bowen, J. E.; Corti, D.; Veesler, D., Structure-guided covalent stabilization of coronavirus spike glycoprotein trimers in the closed conformation. *Nat Struct Mol Biol* **2020,** *27* (10), 942-949.
17. Cai, Y.; Zhang, J.; Xiao, T.; Peng, H.; Sterling, S. M.; Walsh, R. M., Jr.; Rawson, S.; Rits-Volloch, S.; Chen, B., Distinct conformational states of SARS-CoV-2 spike protein. *Science* **2020,** *369* (6511), 1586-1592.
18. Yurkovetskiy, L.; Wang, X.; Pascal, K. E.; Tomkins-Tinch, C.; Nyalile, T. P.; Wang, Y.; Baum, A.; Diehl, W. E.; Dauphin, A.; Carbone, C.; Veinotte, K.; Egri, S. B.; Schaffner, S. F.; Lemieux, J. E.; Munro, J. B.; Rafique, A.; Barve, A.; Sabeti, P. C.; Kyratsous, C. A.; Dudkina, N. V.; Shen, K.; Luban, J., Structural and Functional Analysis of the D614G SARS-CoV-2 Spike Protein Variant. *Cell* **2020,** *183* (3), 739-751 e8.
19. Wrobel, A. G.; Benton, D. J.; Xu, P.; Roustan, C.; Martin, S. R.; Rosenthal, P. B.; Skehel, J. J.; Gamblin, S. J., SARS-CoV-2 and bat RaTG13 spike glycoprotein structures inform on virus evolution and furin-cleavage effects. *Nat Struct Mol Biol* **2020,** *27* (8), 763-767.

20. Xiong, X.; Qu, K.; Ciazynska, K. A.; Hosmillo, M.; Carter, A. P.; Ebrahimi, S.; Ke, Z.; Scheres, S. H. W.; Bergamaschi, L.; Grice, G. L.; Zhang, Y.; Collaboration, C.-N. C.-B.; Nathan, J. A.; Baker, S.; James, L. C.; Baxendale, H. E.; Goodfellow, I.; Doffinger, R.; Briggs, J. A. G., A thermostable, closed SARS-CoV-2 spike protein trimer. *Nat Struct Mol Biol* **2020,** *27* (10), 934-941.
21. Ke, Z.; Oton, J.; Qu, K.; Cortese, M.; Zila, V.; McKeane, L.; Nakane, T.; Zivanov, J.; Neufeldt, C. J.; Cerikan, B.; Lu, J. M.; Peukes, J.; Xiong, X.; Krausslich, H. G.; Scheres, S. H. W.; Bartenschlager, R.; Briggs, J. A. G., Structures and distributions of SARS-CoV-2 spike proteins on intact virions. *Nature* **2020,** *588* (7838), 498-502.
22. Bangaru, S.; Ozorowski, G.; Turner, H. L.; Antanasijevic, A.; Huang, D.; Wang, X.; Torres, J. L.; Diedrich, J. K.; Tian, J. H.; Portnoff, A. D.; Patel, N.; Massare, M. J.; Yates, J. R., 3rd; Nemazee, D.; Paulson, J. C.; Glenn, G.; Smith, G.; Ward, A. B., Structural analysis of full-length SARS-CoV-2 spike protein from an advanced vaccine candidate. *Science* **2020,** *370* (6520), 1089-1094.
23. Chen, W.; Dev, J.; Mezhyrova, J.; Pan, L.; Piai, A.; Chou, J. J., The Unusual Transmembrane Partition of the Hexameric Channel of the Hepatitis C Virus. *Structure* **2018,** *26* (4), 627-634 e4.
